# Dopamine D1 receptor activation and cAMP/PKA signalling mediate Brd4 recruitment to chromatin to regulate gene expression in rat striatal neurons

**DOI:** 10.1101/2021.07.01.450754

**Authors:** Jace Jones-Tabah, Ryan D. Martin, Jennifer J. Chen, Jason C. Tanny, Paul B.S. Clarke, Terence E. Hébert

## Abstract

The activity of striatal medium-spiny projection neurons is regulated by dopamine acting principally at D1 and D2 dopamine receptors. The dopamine D1 receptor (D1R) is a Gα_s/olf_-coupled GPCR which activates a cAMP/PKA/DARPP-32 signalling cascade that increases excitability and facilitates plasticity, partly through the regulation of transcription. Transcriptional regulation downstream of the D1R involves the activation of PKA, which can translocate to the nucleus to phosphorylate various targets. The chromatin reader Brd4 regulates transcription induced by neurotrophic factors in cortical neurons and has also been implicated in dopamine-dependent striatal functions. Brd4 is activated by phosphorylation; this facilitates its binding to acetylated histones at promoters and enhancers. In non-neuronal cells, Brd4 is recruited to chromatin in response to PKA signalling. However, it is unknown whether Brd4 is involved in transcriptional activation by the D1R in striatal neurons. Here, we demonstrate that cAMP/PKA signalling increases Brd4 recruitment to dopamine-induced genes in striatal neurons, and that knockdown or inhibition of Brd4 modulated D1R-induced gene expression. Specifically, inhibition of Brd4 with the bromodomain inhibitor JQ1 suppressed the expression of ∼25% of D1R-upregulated genes, while increasing the expression of a subset of immediate-early genes, including *Fos* and *Jun*. This pro-transcriptional effect of JQ1 was P-TEFb-dependent, and mediated through inhibition of the BD1 bromodomain of Brd4. Finally, we report that JQ1 treatment downregulated expression of many GPCRs and also impaired ERK1/2 signalling in striatal neurons. Our findings identify Brd4 as a novel regulator of D1R-dependent transcription and delineate complex bi-directional effects of bromodomain inhibitors on neuronal transcription.

## Introduction

Dopamine signalling to striatal medium-spiny neurons (MSNs) is critical in controlling movement, processing motivational stimuli, and facilitating learning (1). In the striatum, dopamine signals principally through D1 and D2 dopamine receptors to regulate both the acute excitability of MSNs, and longer lasting forms of synaptic plasticity (2). These latter effects depend on the regulation of nuclear signalling events that modulate gene expression (3, 4). Dysregulation of dopamine-dependent transcription induced by D1 receptor (D1R) signalling also contributes to maladaptive striatal remodeling that occurs in disorders such as drug addiction and L-DOPA-induced dyskinesia (5–7).

All MSNs express dopamine receptors, but the expression of D1 and D2 receptors is largely segregated, with approximately equal proportions of MSNs expressing each receptor (8, 9). The D1R is a G*α*_s/olf_-coupled GPCR which stimulates the activity of adenylyl cyclase (AC) and protein kinase A (PKA). Striatal MSNs produce large and sustained increases in cAMP levels and PKA activity, mediated by the D1R in response to transient pulses of synaptic dopamine (10, 11). D1R signalling in striatal neurons also produces rapid activation of PKA in the nuclear compartment (12). In addition to PKA, other D1R effectors, such as extracellular-regulated kinases (ERK1/2) and dopamine- and cAMP-regulated neuronal phosphoprotein of 32 kilodaltons (DARPP-32), can move into the nucleus where they mediate sustained phosphorylation of nuclear targets and regulate gene expression (13–16).

Striatal D1R activation rapidly induces the expression of immediate-early genes (IEGs), many of which encode transcription factors that can then further modulate gene expression (17). One mechanism by which D1R-dependent signalling can stimulate IEG transcription is through CREB (cAMP response element binding protein) (17, 18), which is activated by PKA and other downstream protein kinases (19–21). Here, however, CREB is only one player among many; D1R signalling converges on numerous transcription factors (17,22–24) and chromatin regulators (25–30) to control gene expression, and the specific mechanisms by which D1R signalling interacts with transcriptional regulators are only partially known. In the present study, we describe a role for bromodomain-containing protein 4 (Brd4) as a mediator of D1R- and cAMP/PKA-dependent transcription in striatal neurons.

Brd4 belongs to the bromodomain and extra-terminal (BET) family of proteins, which consists of Brd2, Brd3, Brd4 and the testis-specific BrdT isoforms (31). BET proteins are characterized by an extraterminal (ET) domain and two tandem bromodomains (BD1 and BD2) which interact with acetylated lysine residues on target proteins, including histones and transcription factors (32, 33). Through its bromodomains, Brd4 acts as a chromatin reader, interacting with acetylated histones in promoter and enhancer regions. Brd4 can also interact with transcription factors (34), and can activate transcription through chromatin-recruitment of the positive transcription elongation factor b (P-TEFb) (35) and the Mediator complex (36). The activity of Brd4 is regulated by phosphorylation of its phosphorylation-dependent interaction domain (PDID). PDID phosphorylation mediated by casein kinases (CK1 & CK2) produces a conformational change that unmasks the bromodomains, thereby facilitating chromatin binding (37, 38).

Recently, Brd4 was identified as a mediator of depolarization- or brain-derived neurotrophic factor (BDNF)-induced transcription in cortical neurons (39, 40), suggesting that Brd4 may play a broad role in activating stimulus-dependent transcription in neurons. Specifically, BDNF promoted chromatin binding of Brd4 through phosphorylation of the PDID by CK2, subsequently leading to transcriptional activation (39).

Brd4 has also been implicated in *striatal* functions (40–44). Systemic administration of JQ1, a small molecule BET inhibitor, was found to attenuate L-DOPA-induced dyskinesia, a common adverse effect in the treatment of Parkinson’s Disease that is associated with elevated D1R signalling in striatal neurons (41). Psychomotor stimulants such as cocaine also elevate dopamine signalling in the striatum, and systemic or intra-striatal Brd4 inhibition attenuates cocaine-associated behaviors such as cocaine induced place-preference and cocaine relapse (42, 43). These studies suggest a role for Brd4 in regulation of transcriptional responses in the striatum.

At present, only casein kinases have been directly shown to phosphorylate the Brd4 PDID *in vivo.* However, protein kinase A (PKA) can phosphorylate the Brd4 PDID when expressed *in vitro* (38). Furthermore, PKA signalling has been shown to regulate Brd4 recruitment to chromatin in cardiomyocytes (45) and pancreatic β cells (46). In pancreatic β cells, cAMP/PKA-dependent activation of CREB stimulated transcription through recruitment of the acetyltransferase CREB binding protein (CBP/p300), leading to histone acetylation and Brd4 recruitment (46). This argues for the possibility that striatal PKA signalling could promote transcription through regulation of Brd4, either through direct phospho-activation or by CREB/CBP mediated histone acetylation.

While most studies indicate that Brd4 recruitment *stimulates* gene expression in neurons, evidence obtained with BET inhibitors suggests Brd4 may also have a repressive role on IEG expression. IEGs are a subset of activity-regulated genes which are rapidly induced by neuronal activity, and often have RNA polymerase II (RNAP2) pre-loaded at promoter-proximal pause sites to facilitate rapid transcription in response to stimuli (47). While brief (e.g. 10-minute) treatment with BET inhibitors has been found to attenuate IEG expression (39), longer periods of exposure (e.g. 2 hours) either failed to alter IEG expression or in some cases increased it (40). This unexplained observation suggests that our understanding how of BET inhibitors affect neuronal transcription remains incomplete.

Given the intersection between evidence supporting the importance of Brd4 in dopamine-dependent striatal effects, and the potential role of PKA in regulating Brd4 recruitment, we hypothesized that Brd4 was a mediator of D1R-dependent transcriptional regulation in striatal neurons. Here, we used primary neonatal rat neuronal cultures to show that Brd4 was recruited to chromatin in response to cAMP/PKA signalling in striatal neurons, and we then used Brd4 knockdown to implicate Brd4 in D1R-dependent transcriptional responses. Next, using a pharmacological approach, we found that the BET inhibitor JQ1 paradoxically increased the expression of a subset of IEGs, including *Fos* and *Jun*, an effect associated with short-term BET inhibition that specifically requires inhibition of the BD1 bromodomain. Lastly, we examined the effect of prolonged treatment with JQ1 on striatal gene expression and function, finding that BET inhibition downregulated the expression of ion channels and GPCRs, leading to impaired responses to multiple neuromodulators.

## Results

### Nuclear PKA is activated by cAMP signalling in striatal neurons

Striatal neurons express a unique complement of signalling proteins that allow them to produce large and sustained increases in cytosolic cAMP/PKA in response to dopamine (10, 11). It has also been reported that in mouse brain slices, stimulation of the D1R rapidly and robustly activates *nuclear* PKA in striatal neurons, and to a much lesser extent in cortical neurons (12). We first sought to establish a system to study this phenomenon using primary cultures of rat striatal or cortical neurons. To monitor nuclear PKA activity in this system we expressed AKAR (a genetically encoded sensor of PKA activity based on Förster resonance energy transfer (FRET) (48)) tagged with a nuclear localization sequence (NLS) (expression shown in Fig. 1A). We stimulated cells with forskolin (FSK), a strong inducer of adenylyl cyclase (AC), and quantified downstream PKA activity (expressed as percent change in FRET, %ΔF/F) over a 30-minute period in thousands of cultured neurons using high-content fluorescence microscopy (49). FSK activated nuclear PKA in both striatal and cortical neurons, as indicated by increased %ΔF/F relative to vehicle treatment (Fig. 1B). Consistent with previous findings (12), activation of nuclear PKA was greater in striatal compared to cortical neurons. Specifically, striatal responses were approximately 3-fold greater, as measured both by the average fluorescence change in the cell population (Fig. 1B and by the number of individual responding neurons (Fig. 1C). We then quantified downstream expression of the IEG products cFos and cJun using immunofluorescence microscopy (Fig. 1D). Despite evoking a stronger PKA response in striatal neurons, FSK induced cFos expression to a similar degree in striatal and cortical neurons, measured as the average fluorescent intensity (Fig. 1E). A related IEG, cJun, was expressed but not detectably induced by FSK in either cell type (Fig. 1F). Overall, these findings suggest that although both cortical and striatal neurons become activated by AC stimulation (as indicated by cFos induction), they engage different signalling responses such as greater activation of nuclear PKA in striatal neurons.

**Figure 1.**
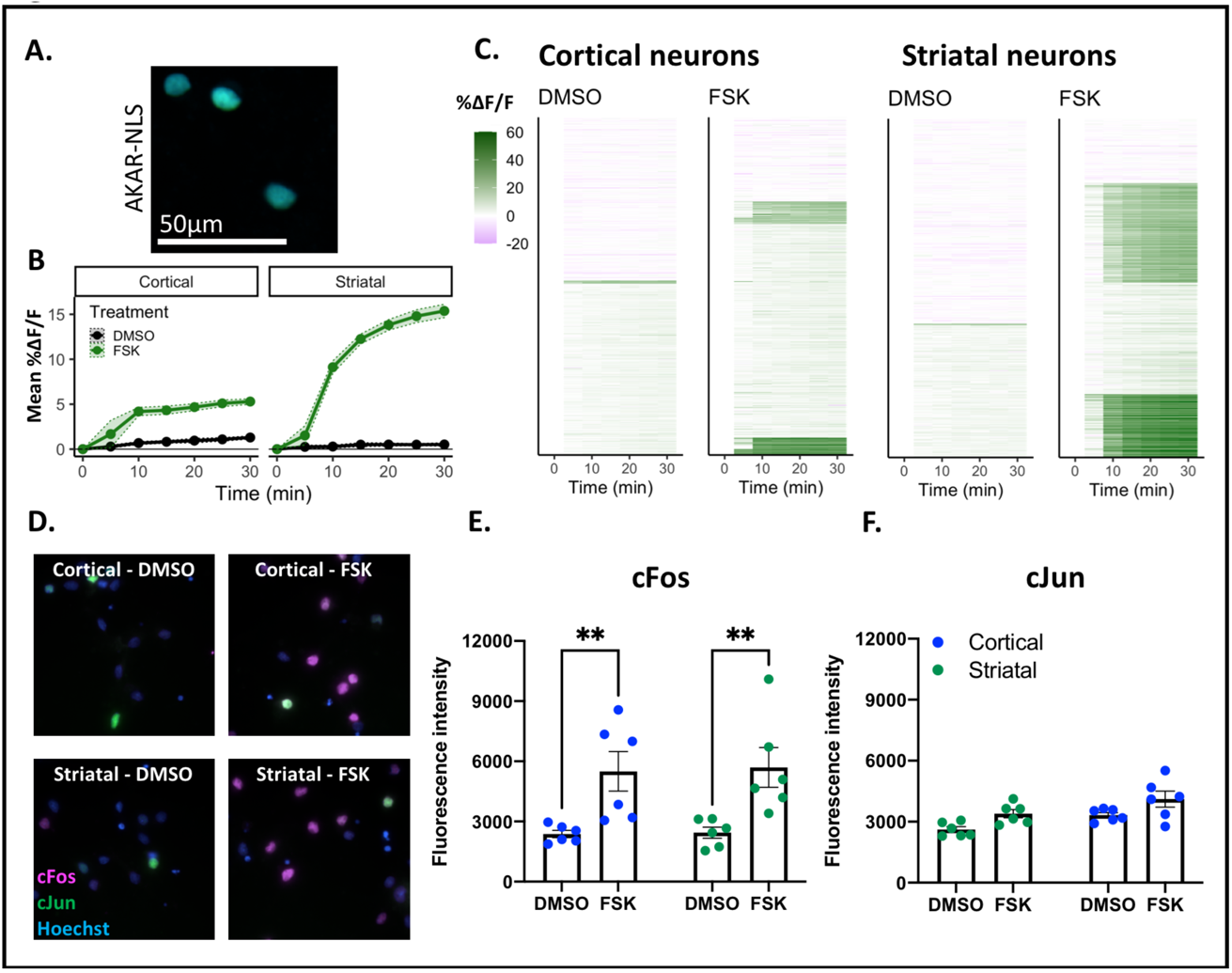
Nuclear PKA is rapidly activated by cAMP signaling in cultured striatal neurons. **A.** Representative image of AKAR-NLS expressed in primary striatal neurons and imaged by high-content microscopy. **B.** Average nuclear PKA response (%ΔF/F AKAR-NLS response) in cortical and striatal cultures stimulated with vehicle (DMSO) or 5 μM forskolin (FSK). Drug was added immediately after acquiring baseline images at time 0. Mean ± SD is shown. **C.** Single-cell breakdown of nuclear PKA responses. Horizontal bars depict individual nuclei and color indicates %ΔF/F relative to baseline. n = 3888 cortical, and 4580 striatal neurons from 5 biological replicates. **D.** Representative image of fixed neurons immunolabelled for cFos (purple) and cJun (green) and with nuclei stained with Hoechst (blue). **E-F.** Quantification of cFos (E) and cJun (F) fluorescence intensity after a 60-minute treatment with vehicle (DMSO) or 5 μM forskolin (FSK). Bars depict mean ± SEM, n = 5 biological replicates with >10,000 nuclei per replicate, ** p<0.01, Bonferroni corrected paired t-tests.

### Cyclic AMP/PKA stimulates Brd4 recruitment to chromatin in striatal, but not cortical neurons

In cortical neurons, BDNF treatment stimulates Brd4 recruitment to the promoter regions of IEGs *Fos*, *Arc* and *Nr4a1* (39). To determine whether PKA activation likewise drives Brd4 recruitment to chromatin, we performed ChIP-qPCR to measure Brd4 chromatin binding in striatal neurons stimulated for 60 minutes with FSK. To this end, we selected the following 8 genes: IEGs (*Fos*, *Fosb*, *Nr4a1, Arc* and *Jun*), secondary response genes (*Drd1, Bdnf*) and a housekeeping gene (*Gapdh*). In each case, we measured Brd4 recruitment either at transcription start-sites (TSS) or in the gene bodies. FSK stimulated Brd4 recruitment to the TSS region of *Fos* and *Fosb*, as well as to the gene-body region of *Fosb*, *Nr4a1* and *Drd1* (Fig. 2). We did not detect FSK-stimulated Brd4 binding to the other loci tested, including at the housekeeping gene *Gapdh*, indicating that the effect of FSK on Brd4 recruitment was gene- or gene location-specific. All genes where Brd4 recruitment was observed were induced by FSK (see Fig. 3C), but some genes where Brd4 recruitment was not detected (e.g. *Arc*, *Bdnf*), were still found to be induced by FSK treatment. We next measured Brd4 chromatin-binding in *cortical* neurons stimulated for 60 minutes with FSK. Brd4 recruitment was not detected at any regions tested in cortical neurons, including loci where significant Brd4 binding was found in striatal neurons (Fig. S1). Hence, the two cell types respond differently to the activation of AC, and our findings support a role for PKA-regulated Brd4 in striatal but not cortical neurons.

**Figure 2.**
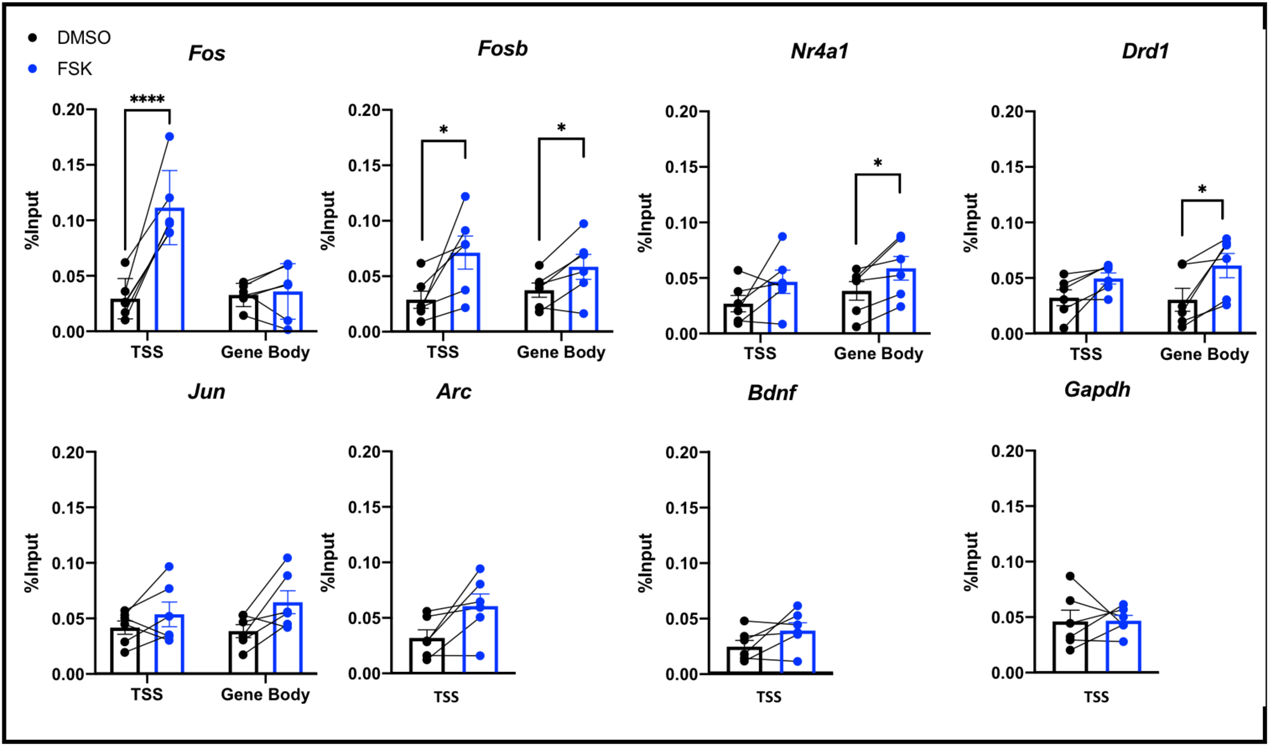
Brd4 is recruited to chromatin in response to cAMP signaling in striatal neurons. ChIP-qPCR showing Brd4 binding to chromatin at the transcription start site (TSS) or gene body for immediate-early genes (*Fos*, *Fosb*, *Arc*, *Jun*, *Nr4a1*), secondary response genes (*Bdnf*, *Drd1*) or a housekeeping gene (*Gapdh*) in striatal neurons treated for 60 minutes with vehicle (DMSO) or 5 μM forskolin (FSK). Values are shown as the percentage input normalized to a spike-in control (see *Experimental Procedures*). Bars depict mean ± SEM, connecting lines indicate biological replicates, n = 6, * p<0.05, **** p<0.0001, Bonferroni corrected paired t-tests.

**Figure 3.**
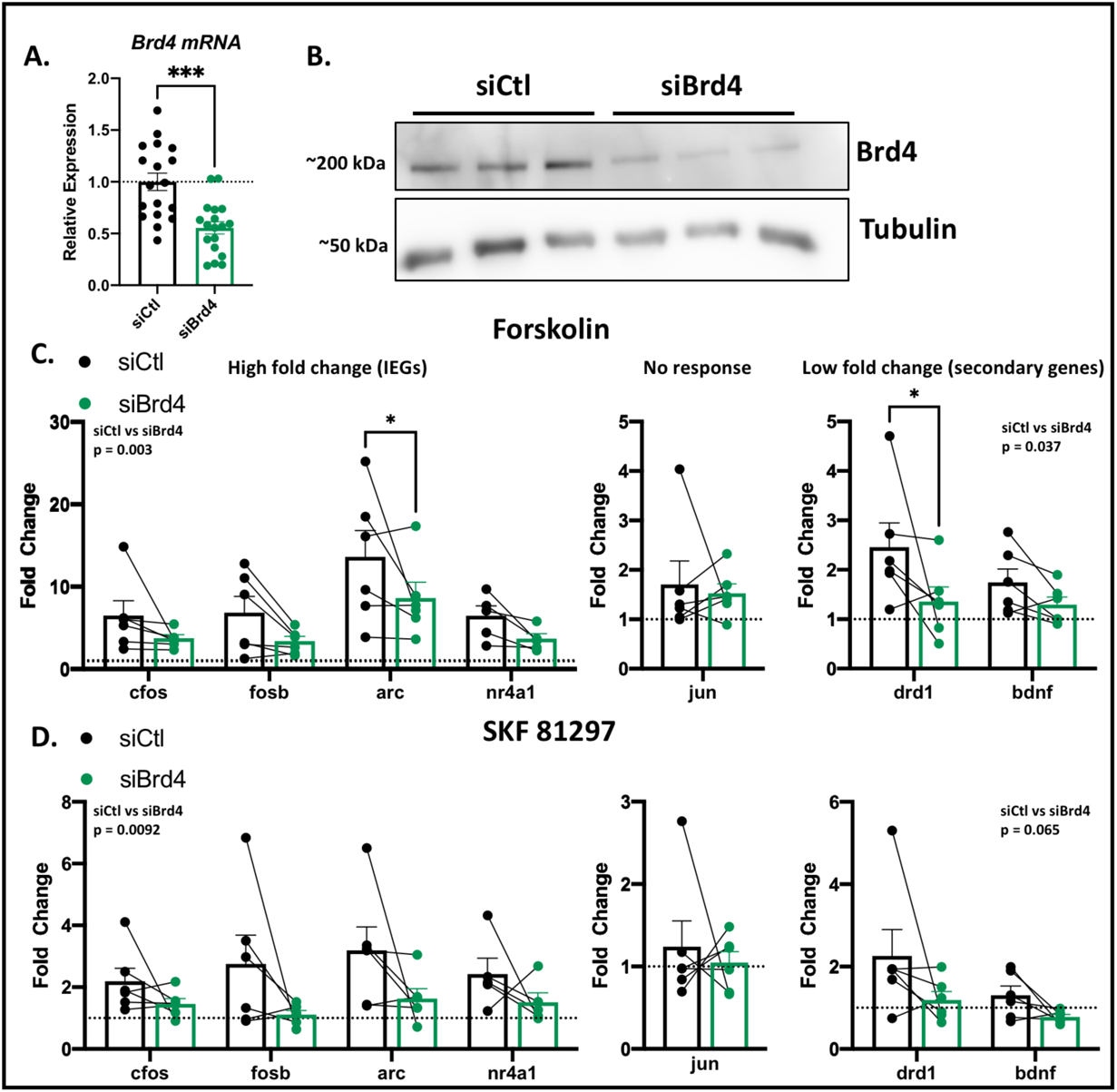
Brd4 knockdown impairs gene expression induced by activation of adenylyl cyclase or D1R in striatal neurons. **A.** Knockdown of Brd4 at 72 hours after siRNA transfection determined by RT-qPCR. Bars depict mean ± SEM, n = 18 biological replicates (representing all vehicle, FSK and SKF 81297 treated samples from panels C and D), independent t-test, ***p<0.001. **B.** Knockdown of Brd4 at 72 hours after siRNA transfection determined by western blot, n = 3 biological replicates. **C-D.** Expression of immediate-early genes (*Fos*, *Fosb*, *Arc*, *Jun*, *Nr4a1*) and secondary response genes (*Bdnf*, *Drd1*) determined by RT-qPCR after a 60-min treatment with 5 μM forskolin (A) or 1 μM of the D1 receptor agonist SKF 81297 (SKF), in striatal neurons transfected with control (siCtl) or Brd4-targetting (siBrd4) siRNA. Values were normalized within each sample to U6 snRNA and are depicted as fold change relative to vehicle treatment for each siRNA condition. Genes were graphed separately based on the degree of response to agonist. Bars depict mean ± SEM, n= 6 biological replicates, 2-way ANOVA p values are indicated for siCtl versus siBrd4 comparisons, * p<0.05 (paired t-test).

### Brd4 is required for D1R- and cAMP-induced gene expression

We next used siRNA-mediated knockdown to determine whether Brd4 contributes to transcriptional activation that is induced either by direct activation of AC or by stimulation of the same pathway via the D1R. Here, we transfected control (siCtl) or Brd4-targeting (siBrd4) siRNAs into cultured striatal neurons, and 72 hours after transfection treated the cells for 60 minutes with vehicle (DMSO), FSK or the D1R agonist SKF 81297 (SKF). Knockdown efficiency at the time of treatment was approximately 50% at the mRNA level (Fig. 3A); in parallel experiments, knockdown at the protein level was confirmed by western blot in three separate experiments (Fig. 3B). We measured expression of the same genes for which Brd4 recruitment was assessed by ChIP in Fig. 2 (with the exception of the housekeeping gene *Gapdh*). For this purpose, gene expression was quantified by RT-qPCR and normalized to levels of the U6 small nuclear RNA (U6 snRNA; Fig 3C-D). Brd4 knockdown alone did not alter baseline expression of any of the genes tested (Fig. S2). However, Brd4 knockdown impaired induction of IEGs and secondary genes by FSK (F [1, 19] = 11.6, p = 0.003, F [1, 10] = 5.71, p = 0.037 respectively; see Fig. 3C) and impaired induction of IEGs by SKF (F [1, 19] = 8.41, p = 0.0092; Fig. 3D). Individual gene comparisons also revealed significant reductions in *Arc* and *Drd1* induction by FSK (Fig. 3C). Consistent with our prior observations, no *Jun* induction was observed with either FSK or SKF treatment. These data indicate that Brd4 contributes to both D1R- and AC-induced expression of IEGs and secondary response genes.

### BET inhibition increases D1R-induced expression of some IEGs

We next focused on the effects of the D1R agonist, and took a pharmacological approach to interrogate the role of Brd4 on gene expression induced by receptor activation. As described previously, small molecule BET bromodomain inhibitors such as JQ1 (50) attenuated BDNF-dependent transcription in cortical neurons through inhibition of Brd4 (39, 40). However, some evidence suggests that certain IEGs, such as *Fos*, are resistant to, or even enhanced by BET inhibition (40). In view of these mixed findings, we investigated the possible effects of BET inhibition in SKF-stimulated striatal neurons. To this end, we treated primary striatal neurons with SKF with or without JQ1, for 60 minutes or 24 hours, and measured the expression of several IEGs by RT-qPCR (Fig. 4, Fig. S3).

**Figure 4.**
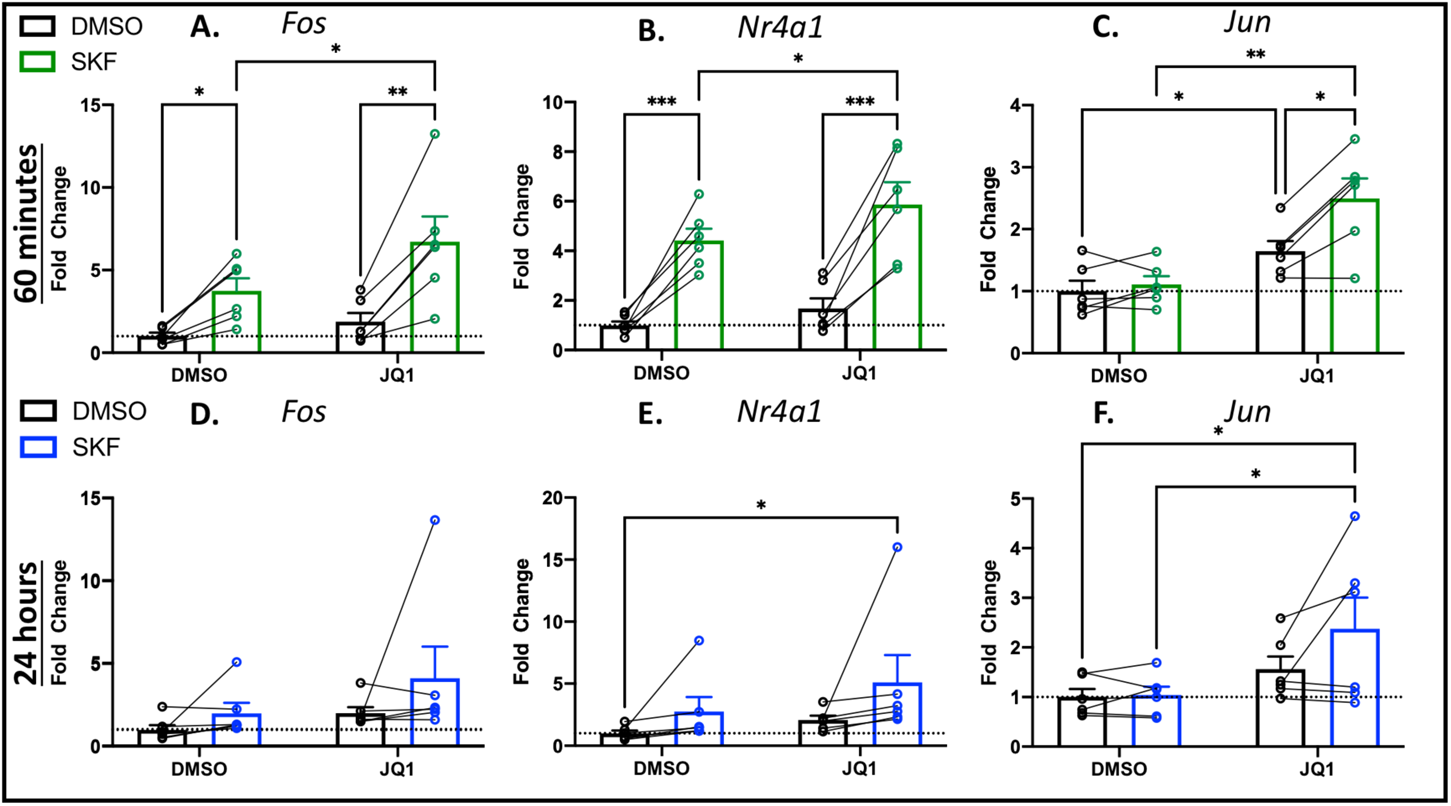
BET inhibition with JQ1 potentiates immediate-early gene expression induced by D1R activation. **A-B.** Expression of three IEGs (*Fos*, *Nr4a1, Jun*) was measured by RT-qPCR either 60 minutes (A) or 24 hours (B) after treatment with vehicle (DMSO) or 1 μM of the D1 receptor agonist SKF 81297 (SKF), with or without 2 μM JQ1. Bars depict mean ± SEM, connecting lines indicate biological replicates, n = 6 biological replicates, * p<0.05, ** p<0.01, *** p<0.001 (Bonferroni corrected paired t-tests, select comparisons shown).

After 60-minute treatment, *Fos* and *Nr4a1* were induced by SKF and further increased in the presence of JQ1 (Fig. 4A,B). As we had previously observed (see above), the D1R agonist given alone had no effect on *Jun*, but unexpectedly JQ1 treatment significantly increased *Jun* expression, and this was further increased when combined with SKF treatment (Fig. 4C). This result suggested that BET inhibition can facilitate induction of IEGs which are not normally coupled to D1R signalling.

After 24-hour treatment, no effects were detected on *Fos* expression (Fig. 4D). In contrast, *Nr4a1* and *Jun* expression were again elevated in striatal neurons treated with combined SKF and JQ1 (Fig. 4E-F), suggesting that induction of these genes may be maintained up to 24 hours after treatment with both SKF and JQ1. Other IEGs tested (*Fosb, Arc* and *Egr1*) were induced by SKF but unaffected by combined treatment with JQ1 (Fig. S3). Together, these data confirm previous reports that the induction of some IEGs is not only resistant to, but is in fact promoted by BET inhibition, and extends this finding to striatal neurons stimulated with a D1R agonist.

### Genome-wide effect of BET inhibition on D1R induced gene expression

To characterize the genome-wide effects of BET inhibition in the context of D1R activation, we next performed RNA sequencing (Fig. 5, Table S1-S3). Consistent with previous findings (17), the effect of the D1R agonist on gene expression was largely stimulatory, and after a 60-min treatment with SKF, we identified 174 and 7 genes that were up- and down-regulated, respectively, in a significant manner (Log_2_ Fold Change > 0.25, and p < 0.01) (Fig. 5A). Treatment for 60 minutes with JQ1 alone significantly up- and down-regulated ∼150 genes in each direction (Fig. 5B light green and purple circles, respectively). When we compared neurons treated with SKF alone to those treated with SKF and JQ1, we identified clusters of 42 and 23 SKF-upregulated genes that were attenuated or increased respectively by co-treatment with JQ1 (Log_2_ Fold Change > 0.25, and p < 0.01 relative to SKF alone) (Fig. 5B blue and yellow circles). Relative expression changes across treatment conditions for these JQ1-attenuated and JQ1-increased genes are shown (Fig. 5C,D). Nearly 70% (16/23) of the genes whose upregulation by SKF was *increased* by addition of JQ1, were also upregulated by JQ1 treatment alone (Fig. 5B). In contrast, only ∼30% (12/42) of the SKF-induced genes that were attenuated by JQ1 showed repression by JQ1 alone.

**Figure 5.**
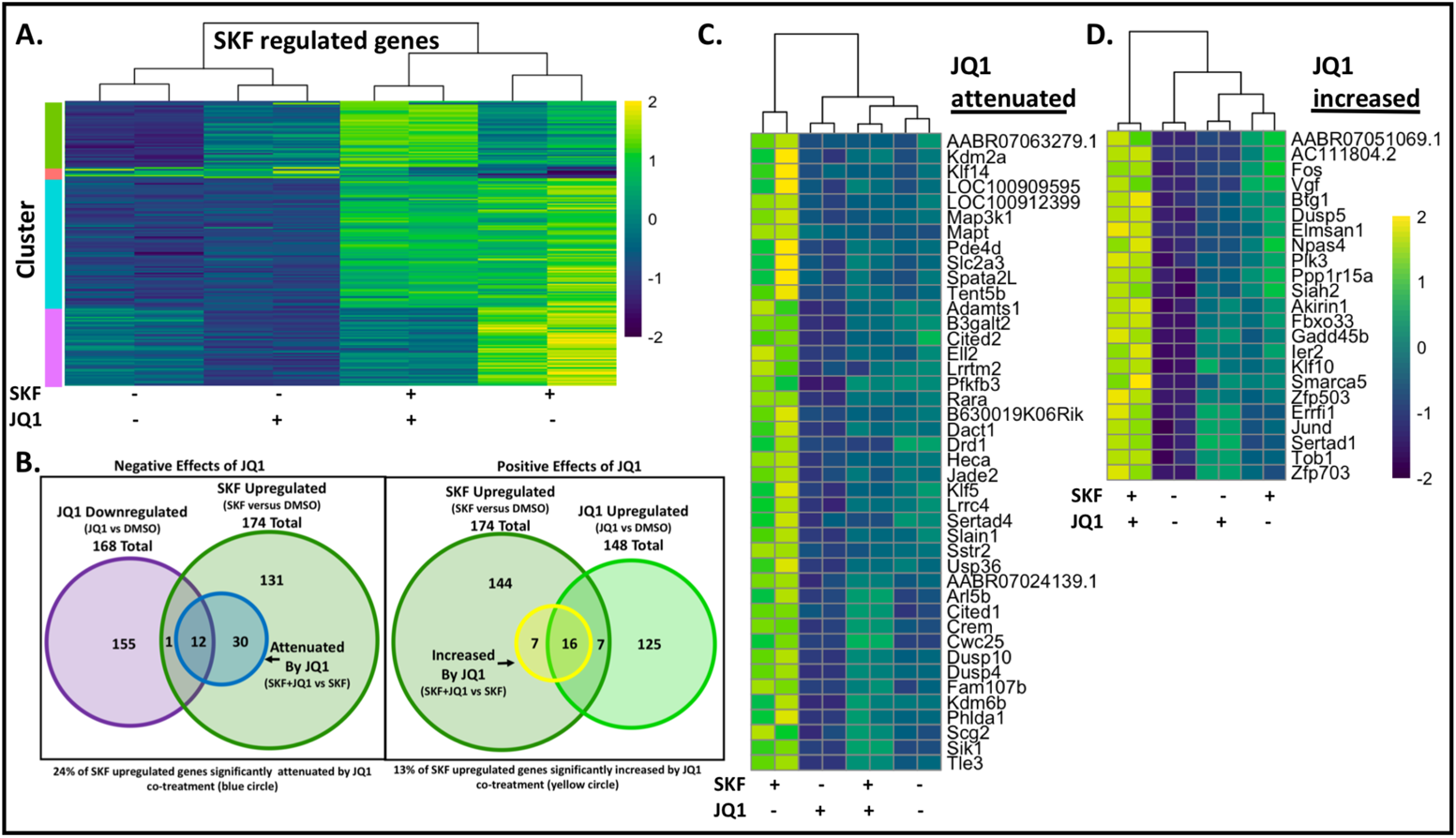
Genome wide effects of BET inhibition on gene expression induced 60 minutes after D1R activation in striatal neurons. **A.** Heat map generated for genes differentially regulated by D1R activation (Log_2_ Fold Change > 0.25, and p < 0.01 for 1 μM SKF 81297 (SKF) relative to vehicle/vehicle condition). Each row depicts the normalized z score for an individual gene across treatment conditions, with value indicated by color. Side-by-side sub-columns represent biological replicates. K-means clustering was performed to identify subsets of genes with distinct patterns of change across treatment conditions. **B.** Venn diagrams depicting SKF 81297-upregulated genes and JQ1 up- or downregulated genes (Log_2_ Fold Change > 0.25, and p < 0.01 relative to vehicle/vehicle condition). Attenuated (blue) and Increased (yellow) groups indicate genes significantly upregulated by SKF, that were significantly increased or attenuated in the presence of JQ1 (Log_2_ Fold Change > 0.25, and p < 0.01 for SKF/JQ1 relative to SKF/vehicle condition). **C-D.** Normalized z-scores of genes upregulated by SKF and attenuated (C) or increased (D) in the presence of JQ1. Side-by-side sub-columns represent biological replicates.

Genes whose induction by SKF was enhanced by JQ1 included numerous IEGs (Fig. 5D), several of which (*Fos*, *Dusp5*, *Fbxo33*, *Gadd45b*, *Ier2*, and *Npas4*) have previously been shown to be regulated in neurons by basal “pre-loading” of RNAP2 at the promoter-proximal pause site (51). In comparison, none of the JQ1-attenuated genes are known to be regulated by basal pausing. Many of these IEGs were also upregulated by JQ1 treatment alone (Fig. 5D), and based on published data, many of the same IEGs were also upregulated in cortical neurons treated for 2 hours with a structurally distinct BET inhibitor, iBET858 (40). Hence, the ability of BET inhibitors to induce the expression of select IEGs appears conserved across multiple neuronal subtypes, and is unlikely to be mediated by off-target effects of a single specific inhibitor.

### JQ1 stimulates P-TEFb dependent transcription

Among the most JQ1-upregulated genes in our RNAseq dataset was *Hexim1* (Fig. 6A) and this gene was also induced in neurons by iBET858 (40). Using RT-qPCR, we found that after a 60-minute treatment, *Hexim1* was upregulated by JQ1, but unaffected by SKF (Fig. 6B). Hexim1 forms part of the 7SK snRNP that sequesters and inactivates P-TEFb. This interaction can be disrupted by Brd4, resulting in both the release and activation of P-TEFb (35, 52). *Hexim1* transcription is stimulated by P-TEFb and forms a negative feedback loop to regulate P-TEFb activity (53). Thus, the increase in *Hexim1* transcription suggested an increase in P-TEFb activity. Therefore, we next asked whether the ability of JQ1 to induce or potentiate IEG expression was P-TEFb dependent. Here, we again used high-content imaging with immunofluorescence to measure cFos and cJun protein expression in primary striatal neurons treated for 60 minutes or 6 hours with vehicle (DMSO) or SKF and the indicated inhibitors (Fig. 6C-F). After a 60-minute treatment, iCdk9, an inhibitor of P-TEFb kinase activity (54) abrogated the increase in cFos or cJun expression induced by either SKF or SKF and JQ1 (Fig. 6C,D). Unexpectedly, 6-hour treatment with iCdk9 significantly *increased* cFos levels, even in the absence of SKF; an apparent attenuation of this iCdk9 effect by JQ1 was not statistically significant (Fig. 6E). In the case of cJun, 6-hour treatment with JQ1 alone or with SKF significantly increased expression, and this was abrogated by iCdk9 (Fig. 6F). Although the mechanism by which iCdk9 increased cFos expression remains unclear, the data presented in Fig. 6 together suggest that the ability of JQ1 to induce IEGs such as *Jun* is generally dependent on P-TEFb kinase activity.

**Figure 6.**
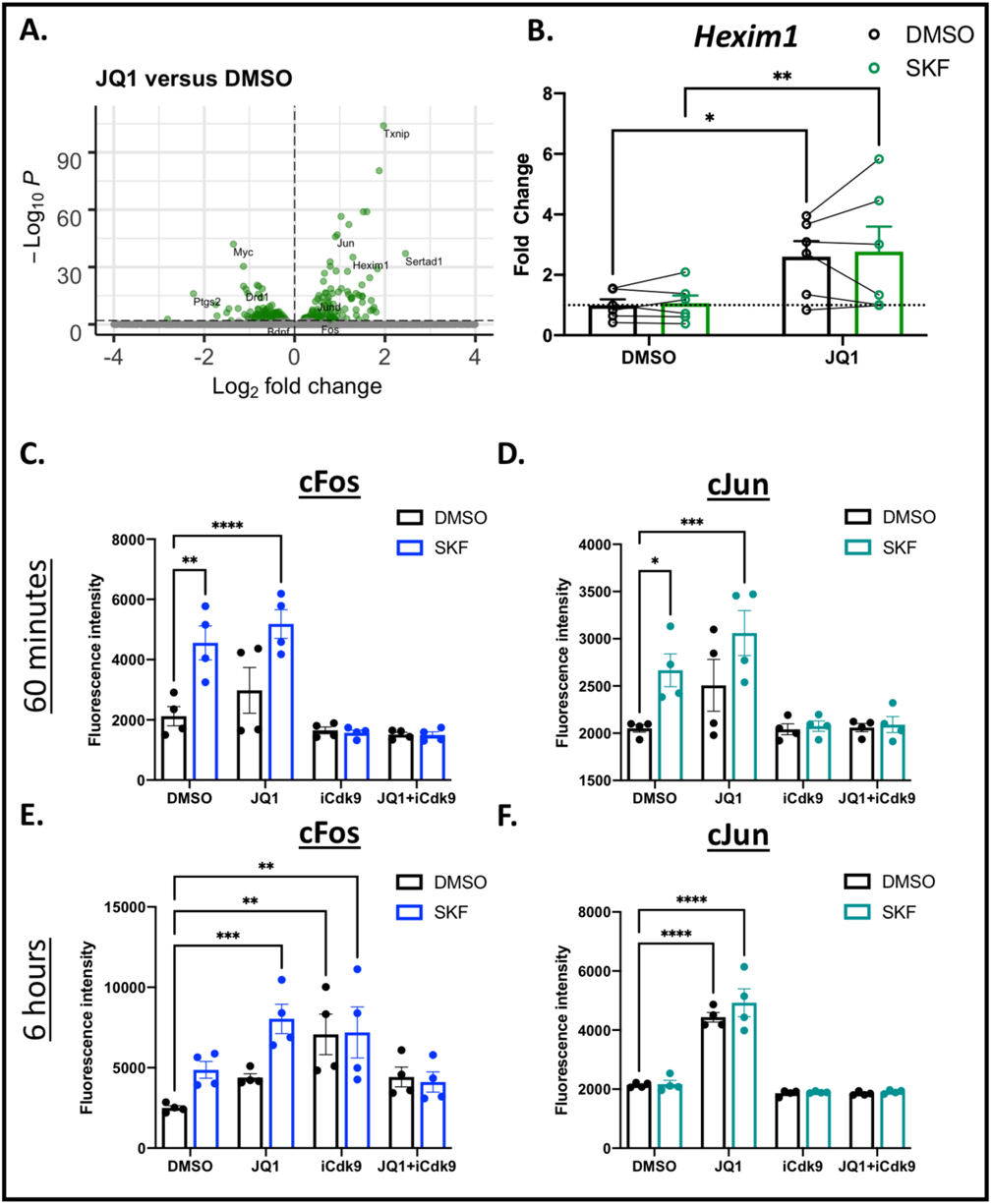
JQ1 induces *Hexim1* and stimulates immediate-early gene transcription through a P-TEFb-dependent mechanism. **A.** Volcano plot of differentially expressed genes following a 60-minute treatment with JQ1 relative to vehicle (DMSO). **B.** Expression changes in *Hexim1* measured by RT-qPCR after 60-minute treatment with SKF 81297 (SKF) and/or JQ1. Bars depict mean ± SEM, connecting lines indicate biological replicates, n = 6 biological replicates, * p<0.05, ** p<0.01 (Bonferroni corrected paired t-tests, select comparisons shown). **C,D.** Quantification of cFos fluorescence intensity after 60 minutes or 6 hours. **E, F.** Quantification of cJun fluorescence intensity after 60 minutes or 6 hours. Bars depict mean ± SEM, n = 4 biological replicates, * p<0.05, ** p<0.01, *** p< 0.001, **** p<0.0001 (Bonferroni corrected t-tests, select comparisons shown).

### Inhibition of BD2 does not stimulate IEG production

When cells are not dividing, the majority of Brd4 is stably bound to chromatin (55, 56). However, this binding is disrupted by JQ1 (39), thereby increasing the amount of available, non-chromatin bound Brd4. Hence, we hypothesized that the ability of JQ1 to stimulate IEGs through P-TEFb was mediated by liberation of Brd4 from stable binding to chromatin. JQ1 inhibits binding of both the Brd4 bromodomains (BD1 and BD2). These bromodomains perform different functions. Notably, BD1 has been shown to mediate the stable association of Brd4 with diacetylated nucleosomes and is generally required for Brd4 association with chromatin, whereas there is an additional requirement for BD2 to mediate certain Brd4 functions such as induction of activity-regulated genes (57).

To interrogate the specific contribution of BD2, we treated striatal neurons for 60 minutes with a BD2-selective inhibitor (iBD2 (57)) alone or in combination with SKF, and again performed RNAseq (Fig. 7A-E, Table S4-S6). Based on previous characterizations of iBD2, the selected dose of 1 μM is expected to produce near-maximal engagement of BD2, without engaging BD1 (57). Unlike JQ1, iBD2 alone had limited effects on gene expression, significantly up- and down-regulating only 22 and 2 genes respectively (Fig. 7A). Likewise, when we examined SKF-regulated genes, we also observed limited effects of iBD2 co-treatment (Fig. 7B). Specifically, only one SKF-induced gene was significantly attenuated (*Drd1*) while three were increased (*Arc*, *Egr5*, *Jun*) by co-treatment with iBD2. The number and composition of genes significantly induced by SKF were largely consistent between the JQ1 and iBD2 RNAseq experiments, suggesting that iBD2’s lack of effect was not related to differences in culture conditions or agonist effect (Fig. 7C).

**Figure 7.**
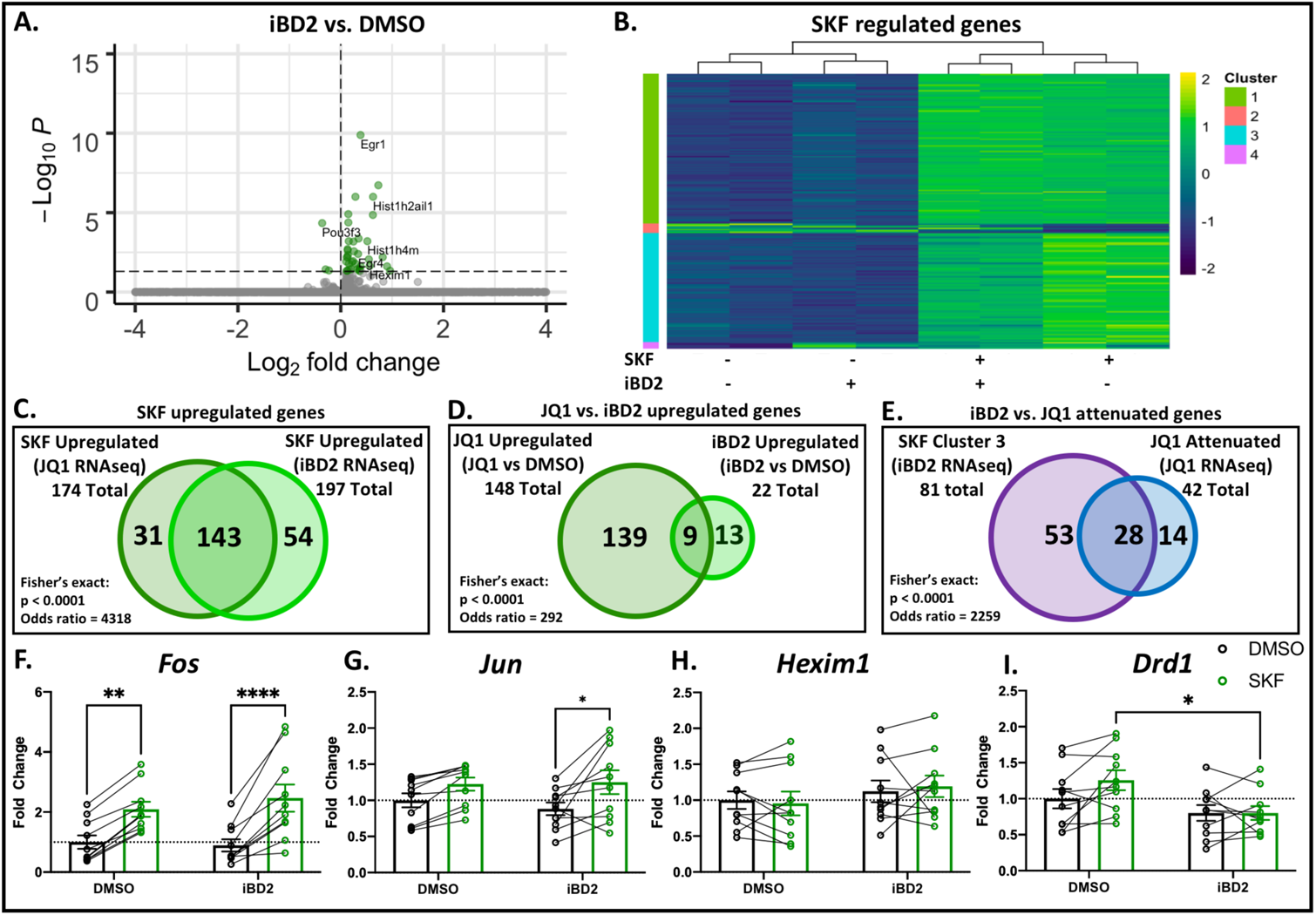
Limited effects of BD2 inhibition on D1R-induced gene expression. **A.** Volcano plot of differentially expressed genes following a 60-minute treatment with iBD2 relative to vehicle (DMSO). **B.** Heat map generated for genes differentially regulated by D1R activation (Log_2_ Fold Change > 0.25, and p < 0.01 for 1 μM SKF 81297 (SKF) relative to vehicle/vehicle condition). Each row depicts the normalized z score for an individual gene across treatment conditions, with value indicated by color. Side-by-side sub-columns represent biological replicates. K-means clustering was performed to identify subsets of genes with distinct patterns of change across treatment conditions **C.** Comparison between the genes upregulated after 60-minute treatment with SKF in the present RNAseq and the previously described RNAseq which was presented in Figure 5. **D.** Comparison between genes significantly upregulated after 60-minute treatment with JQ1 versus 60-minute treatment with iBD2. **E.** Comparison of genes in K-means cluster 3 from panel B versus JQ1-attenuated genes from Figure 5C. Significance of overlap for gene-lists was determined by Fisher’s exact test. **F-I.** Expression of *Fos, Jun*, *Hexim1* and *Drd1* were measured by RT-qPCR 60 minutes after treatment with vehicle (DMSO) or 1 μM SKF 81297 (SKF), with or without 1 μM of the BD2 selective inhibitor iBD2. Bars depict mean ± SEM, n = 6 biological replicates, * p<0.05, ** p<0.01, **** p<0.0001 (Bonferroni corrected paired t-tests).

Genes upregulated by iBD2 partially but significantly overlapped with those upregulated by JQ1 (Fig. 7D) suggesting that although the magnitude of iBD2 effect is small, some effects of JQ1 can be recapitulated by the BD2 inhibitor. Among the genes upregulated by both JQ1 and iBD2 were histone genes, and *Hexim1*, although the fold-induction of *Hexim1* was much lower than observed with JQ1 (Log_2_ fold change of 0.56 for iBD2 versus 1.28 for JQ1).

Although expression of only a single SKF-induced gene was attenuated by iBD2 (*Drd1*, Log_2_ Fold Change > 0.25, and p < 0.01 relative to SKF alone), there was a cluster of modestly downregulated genes which did not meet significance and fold change thresholds (Fig. 7B, Cluster 3). The genes in this cluster significantly overlapped with the previously identified genes attenuated by JQ1 (Fig 7E), again suggesting that iBD2 recapitulates some JQ1 effects, but to a lesser extent. Taken together, these findings show that iBD2 largely does not recapitulate either the positive or negative effects of JQ1 in striatal neurons. Although some similar trends were observed, the magnitude of those effects were considerably smaller for iBD2 compared to JQ1, suggesting that it is likely that BD1 inhibition significantly contributes to both the positive and negative effects of JQ1 in this context.

Given the small effect sizes of iBD2 on several genes, we followed up the observations of our RNAseq experiment by performing RT-qPCR on a larger number of biological replicates to independently measure the expression of genes affected by iBD2 or JQ1. iBD2 did not significantly induce *Fos, Jun* or *Hexim1* transcription (Fig. F-H). However, in the presence of SKF, iBD2 co-treatment did suppress *Drd1* expression (Fig. 7I). We further sought to determine whether the lack of iBD2 impact on IEG expression was an artifact of the dose or timepoint examined. We measured cFos and cJun protein expression by high-content microscopy with immunofluorescence at four timepoints following treatment with FSK and/or iBD2, the latter tested at a range of concentrations. No concentration of iBD2 significantly altered cFos or cJun expression (Fig. S4).

### Prolonged BET inhibition causes widespread transcriptional dysregulation in striatal neurons

While JQ1 may acutely upregulate select IEGs, the overall effect of BET inhibition is largely inhibitory at longer timepoints in cortical neurons (40). To assess the consequences of prolonged BET inhibition in striatal neurons, we performed RNAseq experiments on striatal cultures treated for 24 hours with SKF and/or JQ1 (Fig. 8, Table S7-S9). Although the effect of D1R agonist was minimal at this timepoint, BET inhibition for 24 hours led to widespread changes in the transcriptome, irrespective of concurrent D1R activation (1110 and 2620 significantly up- and down-regulated genes respectively) (Fig. 8A). To identify whether specific cellular pathways were differentially effected, we performed Gene Ontology analysis (58, 59) (Fig. 8B), and gene set enrichment analysis using two reference databases, KEGG (60) (Fig. 8C) and Reactome (61) (Fig. 8D), subsequently focusing on commonalities across the three analyses. Upregulated genes were enriched for ribosome function and chromatin regulation, whereas the most significantly downregulated gene-sets were those relating to transmembrane receptors, including GPCRs and ligand-gated ion channels (Fig 8B-D, Fig. S5).

**Figure 8.**
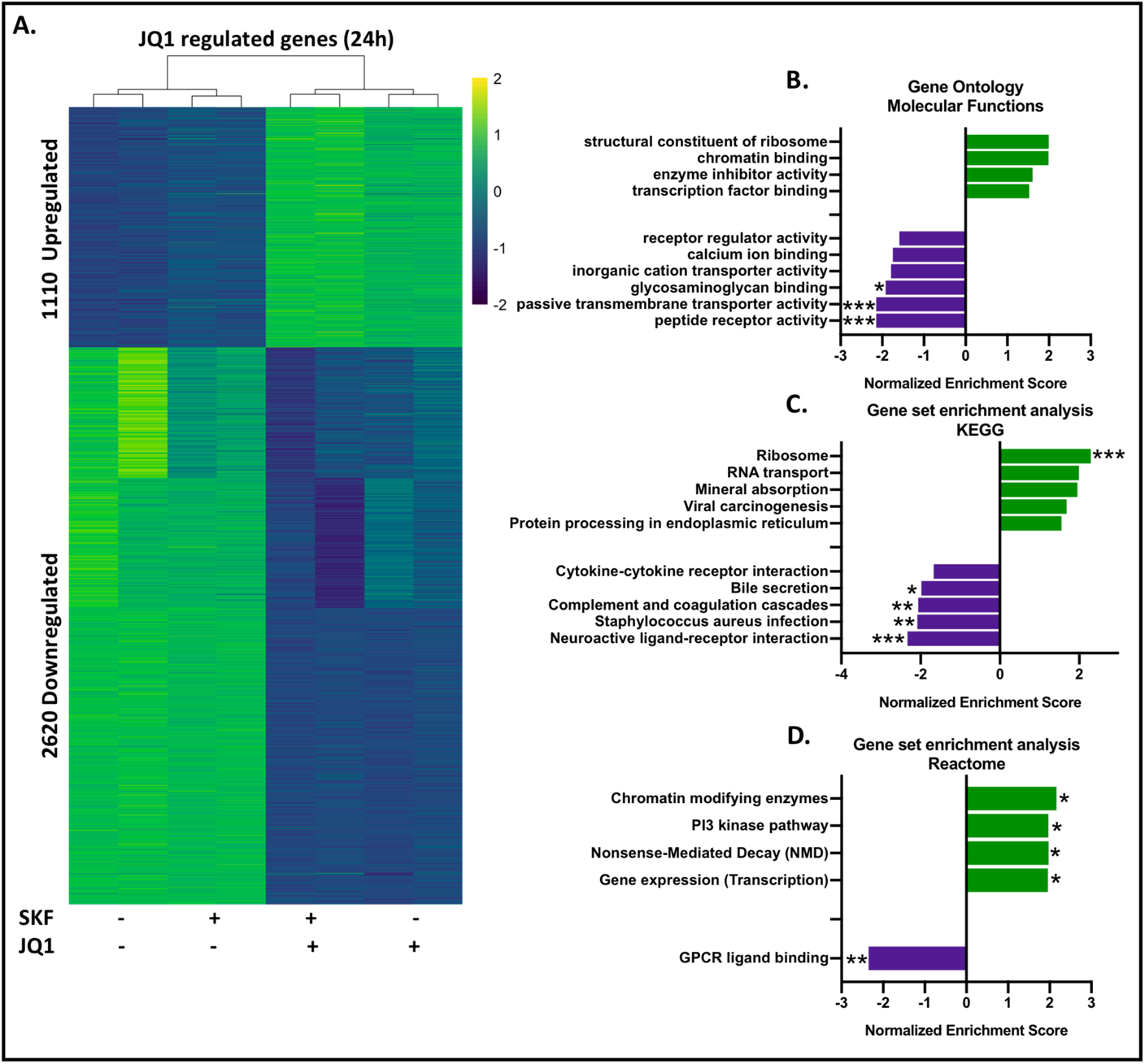
Gene-expression changes in striatal neurons after 24-hour treatment with SKF 81297 and/or JQ1. **A.** Heat map generated for genes differentially regulated by BET inhibition (Log_2_ Fold Change > 0.5, and p < 0.01 for 2 μM JQ1 relative to vehicle/vehicle condition). Each row depicts the normalized z score for an individual gene, with value indicated by color. Side-by-side sub-columns represent biological replicates. **B.** Gene ontology enrichment analysis (Molecular Functions database) for genes differentially regulated by JQ1. **C.** Gene set enrichment analysis (KEGG database) for genes differentially regulated by JQ1. **D.** Gene set enrichment analysis (Reactome database) for genes differentially regulated by JQ1. * FDR < 0.05, ** FDR < 0.01, *** FDR < 0.001.

### Prolonged BET inhibition dysregulates GPCR-dependent signalling in striatal neurons

Since many GPCRs and related genes were downregulated following 24-hour JQ1 treatment (Fig. 8B-D, Fig. S5), we sought to determine whether this transcriptional downregulation would translate into functional changes in GPCR-dependent signalling. We used FRET-based biosensors of PKA (AKAR) and ERK1/2 (EKAR) kinase activity (48) and performed high-content FRET imaging to assess distinct signalling events following stimulation of several receptor types which were downregulated by JQ1 (Table 1). A 24-hour pre-treatment with JQ1 did not detectably alter ligand-induced activation of PKA signalling (Fig. 9A-B, Fig. S6A). In contrast, JQ1 pre-treatment did reduce ERK1/2 signalling induced by specific ligands (treatment x pre-treatment interaction, F [8, 16] = 3.174, p = 0.0235) and this effect was found to be significant specifically when neurons were acutely challenged with dopamine (DA) or serotonin (5HT) (Fig. 9C-D, Fig. S6B). Data for AKAR and EKAR are shown both as the mean %ΔF/F of biological replicates 20 minutes after treatment (Fig. 9B,D) and as mean %ΔF/F of all imaged cells across the full time-course (Fig. S6A-B). We then sought to confirm this reduction in ERK1/2 signalling using an independent methodology: immunofluorescence. Similar to what was observed in the biosensor experiment, ligand-induced increases in phosphorylated (active) ERK1/2 were reduced by JQ1 pre-treatment (treatment x pre-treatment, F [8, 36] = 3.472, p = 0.0046), with significant reductions observed for dopamine (DA), SKF 81297 (SKF) and forskolin (FSK) (Fig. 9E-F). These findings show that treating striatal neurons with JQ1 led to impairments in receptor- and cAMP-dependent activation of ERK1/2.

**Figure 9:**
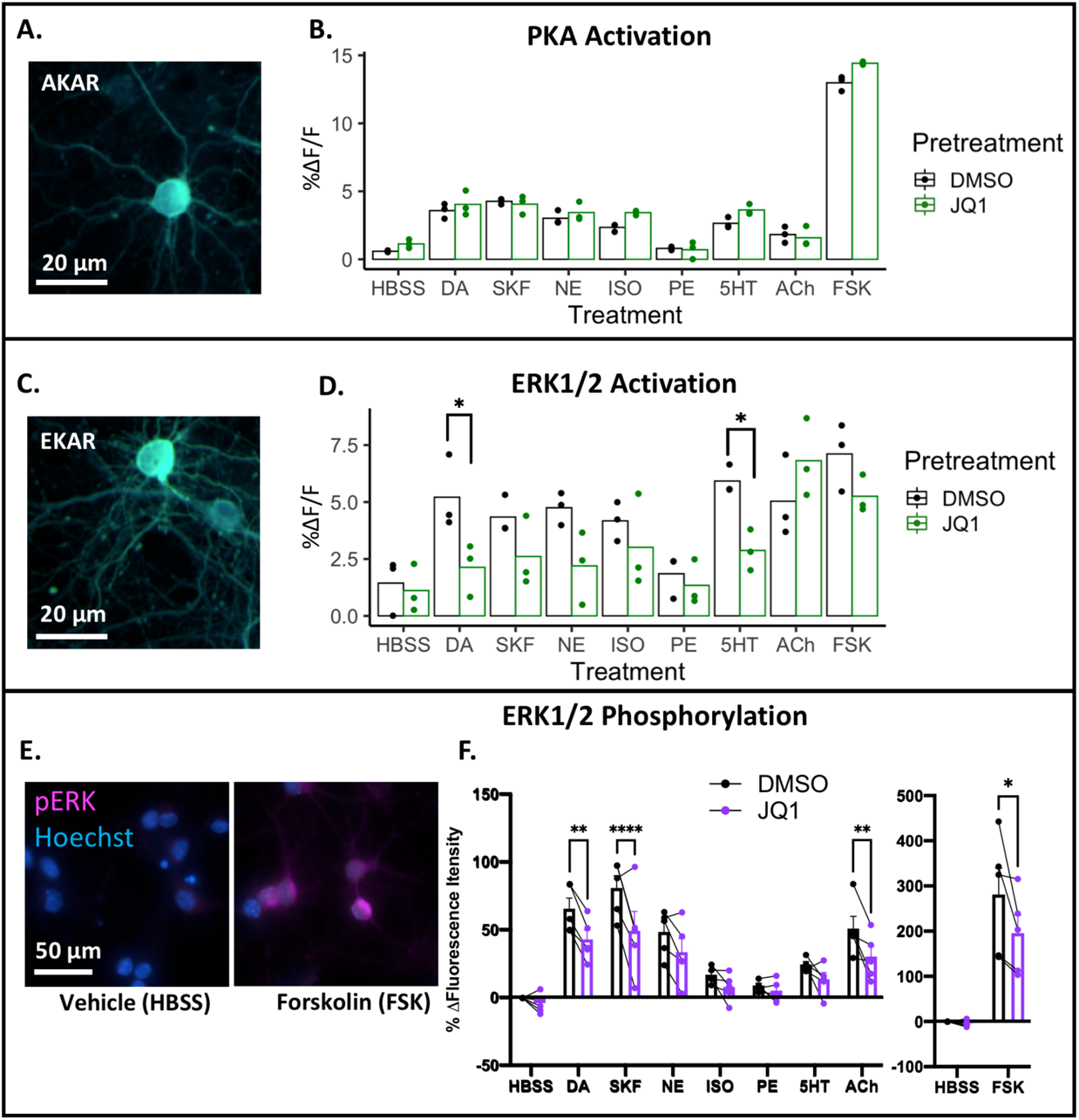
JQ1 attenuates GPCR-dependent ERK1/2 activation. **A.** Representative image depicting the PKA biosensor AKAR expressed in the cytosol of primary striatal neurons. **B.** PKA activation measured as percent change in FRET ratio (%ΔF/F) 20 minutes after treatment with 1 μM of the indicated ligands. **C.** Representative image depicting the ERK1/2 biosensor EKAR expressed in the cytosol of primary striatal neurons. **D.** ERK1/2 activation measured as percent change in FRET ratio (%ΔF/F) 20 minutes after treatment with 1 μM of the indicated ligands. Panels B and D, n = 3 biological replicates, * p<0.05 (Bonferroni corrected paired t-test). **E.** phospho-ERK1/2 immunofluorescence in primary striatal neurons treated for 30 minutes with vehicle (HBSS) or forskolin (FSK). **F.** Quantification of phospho-ERK1/2 immunofluorescence for striatal neurons treated for 30 minutes with 1 μM of the indicated ligands. Values are expressed as percent change from the vehicle/vehicle (DMSO/HBSS) condition. Connecting lines represent individual biological replicates. FSK effects were plotted on a separate scale for visual clarity. In panel F, bars show mean ± SEM, connecting lines show biological replicates, n = 5 biological replicates, * p < 0.05, ** p<0.01, *** p<0.001 (Bonferroni-corrected paired t-tests).

**Table 1.**
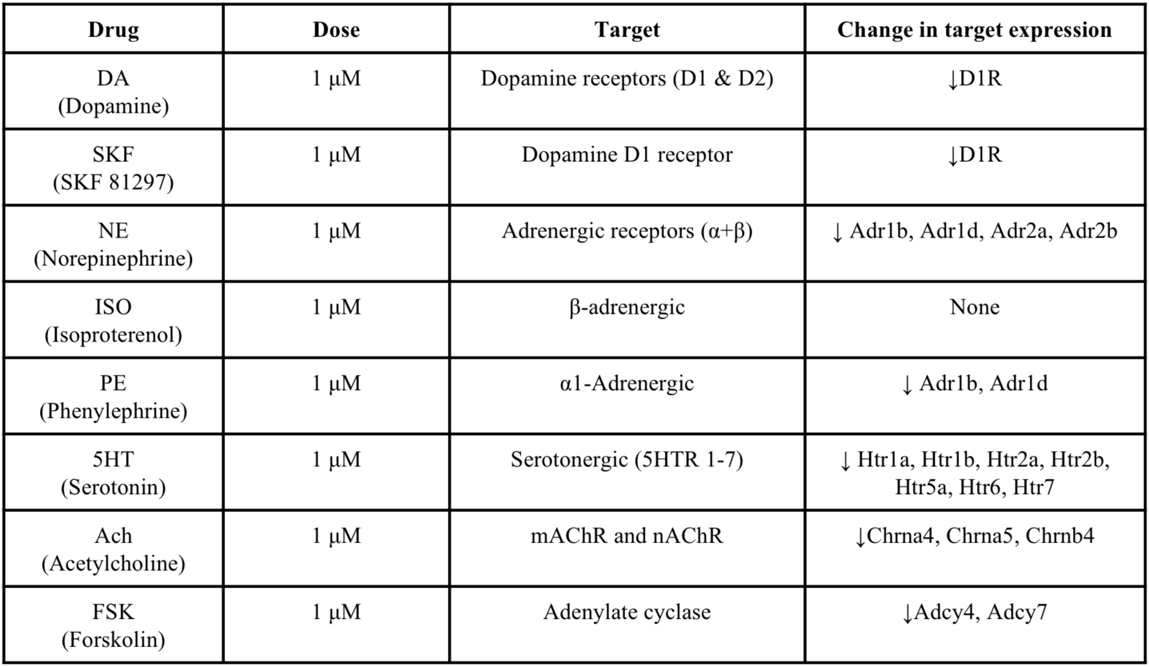

### BET inhibition promotes IEG expression in cortical neurons stimulated with forskolin or BDNF

To date, the role of Brd4 in neurons has been investigated mostly in the context of BDNF-induced gene expression in cortical neurons (39, 40) and here, as described above, the findings with regard to how BET inhibition affects IEG expression are mixed. We therefore investigated whether the ability of JQ1 to promote IEG expression, which we observed in striatal neurons, generalized to cortical neurons. To this end, immunofluorescence and high-content microscopy were used to measure expression of cFos and cJun in primary cortical neurons stimulated with either forskolin or BDNF, in the presence of increasing concentrations of JQ1. JQ1 dose-dependently increased the peak expression of cFos and cJun in cortical neurons challenged with either forskolin or BDNF (Fig. 10A-B). In the absence of JQ1, cFos induction peaked 60-minutes after treatment and then steadily declined, returning to near baseline after 24 hours. In contrast, in the presence of JQ1, the peak cFos induction was shifted from 60 minutes to 6 hours after treatment. We then compared the effects of JQ1 and iBD2 on cortical neuron IEG expression after a longer (i.e. 6-hour) exposure (Fig. 10C-F). JQ1 significantly increased cFos induction by either FSK or BDNF (Fig. 10C); no such effect was seen with iBD2 (Fig. 10E). Similar to our findings in striatal neurons, FSK alone did not induce cJun expression, but co-treatment with JQ1 dose-dependently facilitated induction of cJun by forskolin and BDNF (Fig. 10D,F). In contrast, as we found previously in striatal neurons (Figure 7), iBD2 had no effect (Fig. 10E,F). Taken together, these findings suggest that, JQ1, by inhibiting BD1, stimulates expression of IEGs in both cortical as well as striatal neurons, and can act synergistically with multiple stimuli, including cAMP and receptor tyrosine kinase (BDNF) mediated signalling.

**Figure 10:**
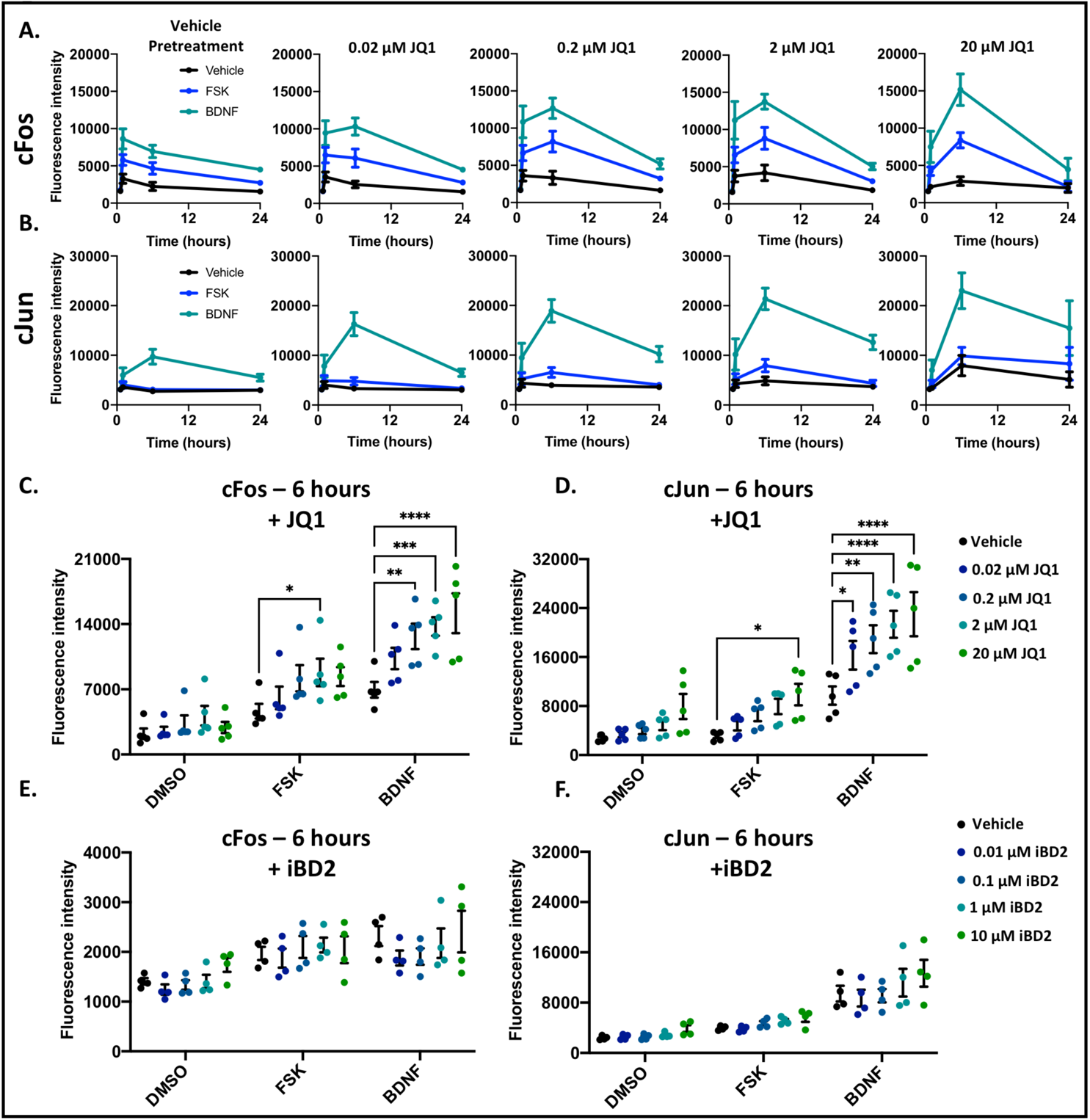
JQ1, but not iBD2, promotes IEG expression in cortical neurons treated with forskolin or BDNF. **A-B.** Time-course of cFos (A) and cJun (B) immunofluorescence in cortical neurons treated with 5 μM forskolin (FSK) or 50 ng/ml brain-derived neurotrophic factor (BDNF) in the presence of JQ1 ranging from 0 to 20 μM. **C.** JQ1 dose-dependent effect on cFos expression after a 6-hour treatment. **D.** JQ1 dose-dependent effect on cJun expression after a 6-hour treatment. **E.** iBD2 dose-dependent effect on cFos expression after a 6-hour treatment. **F.** iBD2 dose-dependent effect on cJun expression after a 6-hour treatment. Bars show mean ± SEM, n = 5 (JQ1) or n = 4 (iBD2) biological replicates, * p<0.05, ** p<0.01, *** p<0.001, **** p<0.0001 (Bonferroni-corrected paired t-tests). *Note:* cFos immunofluorescence was performed using different antibodies for JQ1 experiments versus iBD2 experiments (see methods), and the absolute magnitude of fluorescence intensity should not be compared between these.

## Discussion

The BET protein Brd4 has previously been identified as a mediator of activity-dependent transcription in cortical neurons, specifically downstream of neuronal activity (39, 62) or neurotrophic-factor stimulation (39, 40). In the present study, we have shown that in rat striatal neurons, Brd4 also mediates gene expression downstream of cAMP/PKA dependent signalling induced either directly by AC activation, or by the D1R. We first found that activation of AC with forskolin in striatal neurons led to nuclear PKA activation and Brd4 recruitment to target genes. Forskolin treatment of cortical neurons also activated nuclear PKA (albeit to a lesser extent) and did not detectably increase Brd4 occupancy on chromatin. This latter observation could reflect cell-type differences in the ability of cAMP signals to regulate Brd4, although it is also possible that, given the lower number of responding cortical neurons, Brd4 recruitment was below the limit of detection in our ChIP assay. Although knockdown of Brd4 impaired transcription induced by AC or D1R activation, BET inhibition with JQ1 had bi-directional effects on transcription; notably, a subset of genes showed attenuated expression, as expected, whereas the expression of other genes, largely IEGs, were enhanced in presence of the BRD4 inhibitor. The ability of JQ1 to stimulate IEG expression required P-TEFb activity and was not recapitulated by a selective inhibitor of BD2, suggesting that blockade of BD1 is responsible for this effect. The negative effects of JQ1 on D1R-induced gene expression were only partly recapitulated by BD2 inhibition, indicating that the attenuation of SKF-induced gene expression by JQ1 also involves BD1 inhibition. Prolonged BET inhibition in striatal neurons also broadly downregulated genes involved in shaping neurotransmitter signalling, and impaired GPCR-dependent ERK1/2 activation.

In cortical neurons, as well as in some cancer cell lines, Brd4 is activated through phosphorylation of its PDID (32, 38) and then recruited to acetylated histones, mostly at promoter and enhancer regions. Brd4 phospho-activation is known to be mediated by CK2 (38), and in cortical neurons BDNF promotes Brd4 activation via CK2-dependent phosphorylation (39). In an *in vitro* system, the Brd4 PDID was also phosphorylated by PKA (38). Furthermore, previous studies in cardiomyocytes (45) and pancreatic β cells (46) have linked PKA signalling to Brd4 chromatin-recruitment and transcriptional activation. Collectively, these findings as well as the data presented here indicate that PKA-dependent signalling can activate Brd4, but it remains to be determined whether PKA directly targets Brd4 *in cellulo*.

While it is possible that PKA signalling could promote Brd4 activation through regulation of CK2, we consider this to be unlikely in our experimental system. CK2 is not a known effector of D1R-dependent signalling in striatal neurons. In fact, CK2 has been shown to directly antagonize D1R and G*α*_s_ signalling (63), and also promotes nuclear export of DARPP-32 (15), a key D1R effector that amplifies PKA signalling. Inhibition or knockdown of CK2 amplifies D1R-dependent cAMP signalling (63), and promotes increased nuclear accumulation of DARPP-32 (15). This negative regulation of D1R signalling by CK2 also precluded the use of CK2 inhibitors to assess the contribution of CK2 to Brd4 activation in response cAMP/PKA signalling in striatal neurons. At present, we can only conclude that a PKA-dependent pathway promotes Brd4 activation, and future studies will be required to elucidate the specific mechanisms.

BDNF, in addition to stimulating Brd4 phosphorylation, promotes H3K14 and H4K16 acetylation at IEG promoters, creating binding sites for Brd4 recruitment (39). Electrical stimulation of cortical neurons also promotes rapid histone acetylation at IEG promoter and enhancer regions, leading to Brd4 recruitment and transcriptional activation (62). These findings suggest that stimulus-induced histone acetylation also contributes to Brd4 recruitment. In β-cells, stimulation with forskolin promotes CREB and CREB binding protein (CBP)/p300 recruitment to target genes, leading to CBP-dependent histone acetylation and subsequent Brd4 recruitment (46) suggesting that a similar mechanism operates in striatal neurons.

D1R signalling in striatal neurons has also been linked to histone acetylation and the deposition of other histone modifications which could promote Brd4 recruitment. For example, D1R agonists have been found to promote the adjacent deposition of H3K14ac and H3S10 phosphorylation (H3S10pK14ac) in a PKA-dependent manner (29). Previously-described mechanisms provide a potential link between these modifications and Brd4 recruitment. For example, in a non-neuronal cell line, H3S10 phosphorylation of acetylated H3 residues (in this case H3S10pK9ac) led to recruitment of the adaptor protein 14-3-3 and the histone acetyltransferase MOF, which subsequently acetylated the neighboring histone H4 on K16, and led to Brd4 recruitment (64). An analogous mechanism could link D1R-dependent signalling to Brd4 recruitment, but further studies are required to test this.

The observation that JQ1, but not iBD2, promoted expression of IEGs led us to conclude that inhibition of BD1 function was responsible for the observed gene induction by JQ1 treatment. Based on previous findings, BD1 is thought to mediate the stable binding of Brd4 to chromatin, whereas BD2 is primarily involved in dynamic recruitment to activity-regulated genes (57). Many of the IEGs induced by JQ1 are pre-loaded with poised RNAP2 under basal conditions (51) and such genes are expected to be particularly sensitive to changes in P-TEFb activity, which can release RNAP2 from its paused state. Across a range of cell lines, nearly all Brd4 is chromatin-bound during interphase (55), and this appears to be true for post-mitotic neurons as well (39). We consider it likely that JQ1 stimulates IEG transcription through BD1 inhibition, leading to release of Brd4 from chromatin, which allows for Brd4 to interact with and activate P-TEFb (65). A precedent for this mechanism has been described in non-neuronal cell lines. For example, in immortalized T lymphocytes, JQ1 stimulated the release of P-TEFb from inhibitory interactions with the 7SK snRNP, resulting in assembly of a Brd4-P-TEFb complex, and stimulation of *Hexim1* transcription (66). In a second example in osteosarcoma cells, JQ1 was likewise able to promote release of P-TEFb from the 7SK snRNP, in this case resulting in a global increase in RNA production (67).

Stimulus-dependent release of Brd4 from sequestration on hyperacetylated chromatin may also be involved in activation of Brd4. For example, in response to cellular stress, Brd4 is released from chromatin via nucleosome de-acetylation, allowing Brd4 to interact with and activate P-TEFb (55). Stimulation with UV light or other cellular stressors led to global reduction in Brd4-chromatin occupancy, coupled with an increase in P-TEFb chromatin recruitment, which was blocked in the presence of an HDAC inhibitor. Efficient transcriptional responses to interferon-*α* were also dependent on HDAC-mediated release of Brd4 from hyper-acetylated chromatin (68). This stimulus-dependent release from chromatin depends on both HDACs and H3S10 dephosphorylation mediated by PP1 protein phosphatase (69).

An important limitation to many studies utilizing BET inhibitors, including our own work, is the non-specificity of these drugs across the BET family. Although the importance of Brd4 has been established, using specific genetic techniques such as targeted knockdown or induced degradation, pharmacological inhibition does not strictly phenocopy protein knockdown, as exemplified by the data presented here. The effects of BET inhibitors in the literature are often attributed to inhibition of Brd4, but antagonism of Brd2 and Brd3 cannot be discounted. Unlike Brd4, Brd2 and Brd3 do not interact with P-TEFb, but have been shown to act as histone chaperones, stimulating transcription through the regulation of chromatin (70). In neurons, little is known about the function or importance of these other BET family members, but under some conditions, it has been shown that all three BET proteins work in tandem to regulate transcription. For example, interferon-*γ* stimulated gene expression can be blocked either by non-specific BET inhibitors, or by specific BD2 inhibitors, but targeted degradation of individual BET proteins only partially recapitulated this inhibition (57).

Until recently, most BET inhibitors were also non-selective for the individual bromodomains, BD1 and BD2. The recent development of selective inhibitors has allowed researchers to link the anti-cancer effects of BET inhibitors to BD1, and stimulus-dependent recruitment to BD2 (57). Our data indicate that BD1 and BD2 also play distinct roles in neurons. Specifically, our findings indicate that the majority of the effects of JQ1 that we observed, both on basal and stimulus-induced gene expression, involve BD1, with a lesser contribution of BD2. The future application of more specific pharmacological tools may help clarify the complex and sometimes contradictory observations described for BET function in the brain.

Brd4 inhibitors are being pursued for clinical application, most notably for cancer treatment. For example, several brain-penetrant BET inhibitors are undergoing clinical evaluation for treatment of brain cancers such as glioma, which often overexpress Brd4 (71). Preclinical studies have also suggested that Brd4 inhibitors may have utility in other conditions including cardiovascular and inflammatory diseases (31). Understanding how Brd4 is regulated by different intracellular signalling pathways and in different brain regions could shed light on the potential and pitfalls associated with these drugs.

## Experimental Procedures

### Drugs and Reagents

Unless otherwise noted, reagents were purchased from Sigma-Aldrich. SKF 81297 hydrobromide (Toronto Research Chemicals), forskolin, JQ1 (Selleckchem), iCdk9 (ProbeChem) and iBD2 (also called GSK046, MedChemExpress) stock solutions were prepared in DMSO and stored at −80°C. Potassium chloride (KCl), quisqualic acid (QA – Tocris), and N-methyl-D-aspartic acid (NMDA) were dissolved in Hank’s balanced salt solution (HBSS – Wisent).

### Animals

Sprague-Dawley rat dams with postnatal day 0 pups were purchased from Charles River, Saint-Constant QC, Canada. Animals were maintained on a 12/12-hour light/dark cycle, with free access to food and water. All procedures were approved by the McGill University Animal Care Committee, in accordance with Canadian Council on Animal Care Guidelines.

### Isolation and culture of primary striatal and cortical neurons

Primary cortical and striatal neurons were prepared from freshly dissected Sprague-Dawley rat pups at postnatal day 0. The day before dissection, culture plates were coated with 0.1 mg/ml poly-D-lysine overnight. All solutions were prepared and stored at 4°C. Prior to dissection, plates were washed three times with sterile water and allowed to dry, and solutions were warmed to 37°C. The rat pups were decapitated on ice, and brains were rapidly removed and placed in ice cold Hank’s balanced salt solution (HBSS) without calcium and magnesium (Wisent). The frontal cortex and striatum were dissected and digested separately for 18 min while rotating at 37°C with papain diluted to a final concentration of 20 units/ml in a neuronal medium (Hibernate A Minus Calcium, BrainBits). Next, in order to stop digestion, tissue was transferred for 2 min to HBSS containing 10% fetal bovine serum, 12 mM MgSO_4_, and 10 units/ml DNase1 (Roche). Tissue was pelleted by centrifugation at 300 X *g* for 5 min and supernatant was removed. Tissue was then triturated in HBSS with 12 mM MgSO_4_ and 10 units/ml DNase1 using a fire-polished Pasteur pipette. Once a cell suspension was produced (approximately 20 up and down triturations), it was passed through a 40 µm mesh filter (Fisher) to remove undigested tissue and then centrifuged on an OptiPrep^TM^ gradient as previously described (72), in order to remove cell debris. Purified neurons were then counted and diluted in Neurobasal™-A Medium (NBA) with 1X final concentration of B27 supplement (Gibco), 1% GlutaMAX (Gibco) and 1% penicillin/streptomycin (henceforth referred to as complete NBA) supplemented with 10% fetal bovine serum. Cells were plated at the densities indicated for each type of experiment. Sixteen hours after plating, cells were washed with warm HBSS and media was changed for complete NBA containing 5 µM cytosine-D-arabinoside to inhibit glial cell growth. Cultures were maintained in complete NBA, and media was refreshed by exchanging 30% of the volume with fresh media every 3 days.

### siRNA Transfection

For small interfering RNA (siRNA) transfection, primary neurons were plated on 6-well culture dishes (ThermoFisher) at a density of 2 million cells per well and cultured for 3 days prior to transfection. For each well, 5 μl of 20 μM siRNA (Rat Brd4 SMARTpool, or rat non-targeting control siRNA #5, Dharmacon) was combined with 2.5 μl of Lipofectamine 2000 (Invitrogen) in 100 μl of OptiMEM (Gibco). This mixture was allowed to incubate at room temperature for 20 minutes and then added to cell culture media. Neurons were then cultured as normal until treatments were applied.

### AAV Transduction and high-content FRET microscopy

For FRET imaging experiments, primary neurons were plated on 96-well optical bottom imaging plates (Nunc) at a density of 50,000 cells per well. The FRET-based protein kinase biosensor AKAR3-EV was generously provided by Dr. Michiyuki Matsuda (48) and expressed with a nuclear localization (NLS) peptide sequence using a neuron-specific adeno-associated virus plasmid, pAAV-SynTetOFF (73), kindly provided by Dr. Hiroyuki Hioki. All experiments were performed with AAV serotype 1 produced by the Neurophotonics Platform Viral Vector Core at Laval University, Quebec. Primary neurons were transduced by adding AAV directly into the culture media three days after cell plating, using a multiplicity of infection of 5000 viral genomes/cell. Neurons were then maintained as described above for 7 days prior to imaging.

Live-cell FRET imaging and analysis was performed as we have previously described (49) using an Opera Phenix™ high content confocal microscopy system (Perkin Elmer). Images were acquired using a 40X water-immersion objective using a 425 nm laser for excitation of CFP. Emissions were detected with filters at 435-515 nm (CFP) and 500-550 nm (YFP). Baseline images were acquired and then vehicle or drug solution was added directly to each well, and images were then acquired at the indicated time intervals. FRET ratio was calculated as YFP/CFP and expressed as the percent change in FRET ratio for each cell relative to baseline (%ΔF/F). Following each imaging session, cells were fixed and processed for immunofluorescence, as described in the following section. All image analysis was performed using Columbus™ analysis software (Perkin Elmer) using the a workflow for single cell analysis as previously described (49).

### Cell fixation, immunofluorescence and high-content imaging

For immunofluorescence imaging, experiments primary neurons were plated on 96-well optical bottom imaging plates (Nunc) at a density of 50,000 cells per well. To fix cells, a solution of 4% paraformaldehyde prepared in phosphate-buffered saline (PBS) was added directly into the culture media to reach a final concentration of 2% paraformaldehyde. Fixed cells were washed twice with PBS and permeabilized for 10 minutes using 0.3% Triton in PBS. Blocking was performed for 3 hours at 4°C with 5% bovine serum albumin (BSA) in PBS and then cells were incubated overnight at 4°C with primary antibodies diluted in PBS with 5% BSA. Cells were washed twice with PBS and then incubated for 3 hours at room temperature with secondary antibodies diluted in PBS with 5% BSA. Cells were then washed twice with PBS and incubated in Hoechst dye (Invitrogen) diluted 1:10,000 in PBS. The following primary antibodies and dilutions were used. Anti-cFos polyclonal (1:2000, catalogue #sc-52, Lot #C1010, Santa Cruz), anti-cFos monoclonal (1:5000, mAb #2250 Cell Signalling Technology), and anti-cJun (1:1000, #610326 BD Bioscience), phospho-p44/42 MAPK (Erk1/2) (Thr202/Tyr204) (1:1000, #9101 Cell Signalling Technology). Secondary antibodies were anti-mouse Alexa 488 (1:500; A21236) and anti-rabbit Alexa 647 (1;500; A21245), both from Invitrogen. Fluorescence imaging of fixed cells was performed in the Opera Phenix™ high-content confocal microscope. All image analysis was performed using Columbus™ analysis software (Perkin Elmer). *Note:* cFos antibody from Cell Signalling Technology was used in iBD2 experiments, whereas the cFos antibody from Santa Cruz was used in all other experiments.

### RNA extraction and RT-qPCR

For RT-qPCR experiments, primary neurons were plated on 6-well culture dishes (ThermoFisher) at a density of 2 million cells per well and cultured for 7 days prior to drug treatments. Following the indicated treatments, cells were lysed in TRIzol™ reagent (Invitrogen) and RNA was extracted following the manufacturer’s protocol. Reverse transcription was performed on 1 μg of RNA with random hexamer primers using MMLV-RT enzyme (Promega) according to the manufacturer’s protocol. Gene expression was determined by qPCR using Brightgreen Dye qPCR master mix on a ViiA 7 Real Time PCR System (Applied Biosystems). Primer sequences were designed using NCBI Primer Blast. Primers were supplied by IDT and validated for specificity and efficiency before use. Primer sequences for RT-qPCR are presented in Table 2. Ct values were normalized to U6 snRNA and fold-changes over respective control were calculated using the 2^−ΔΔCt^ method.

**Table 2.**
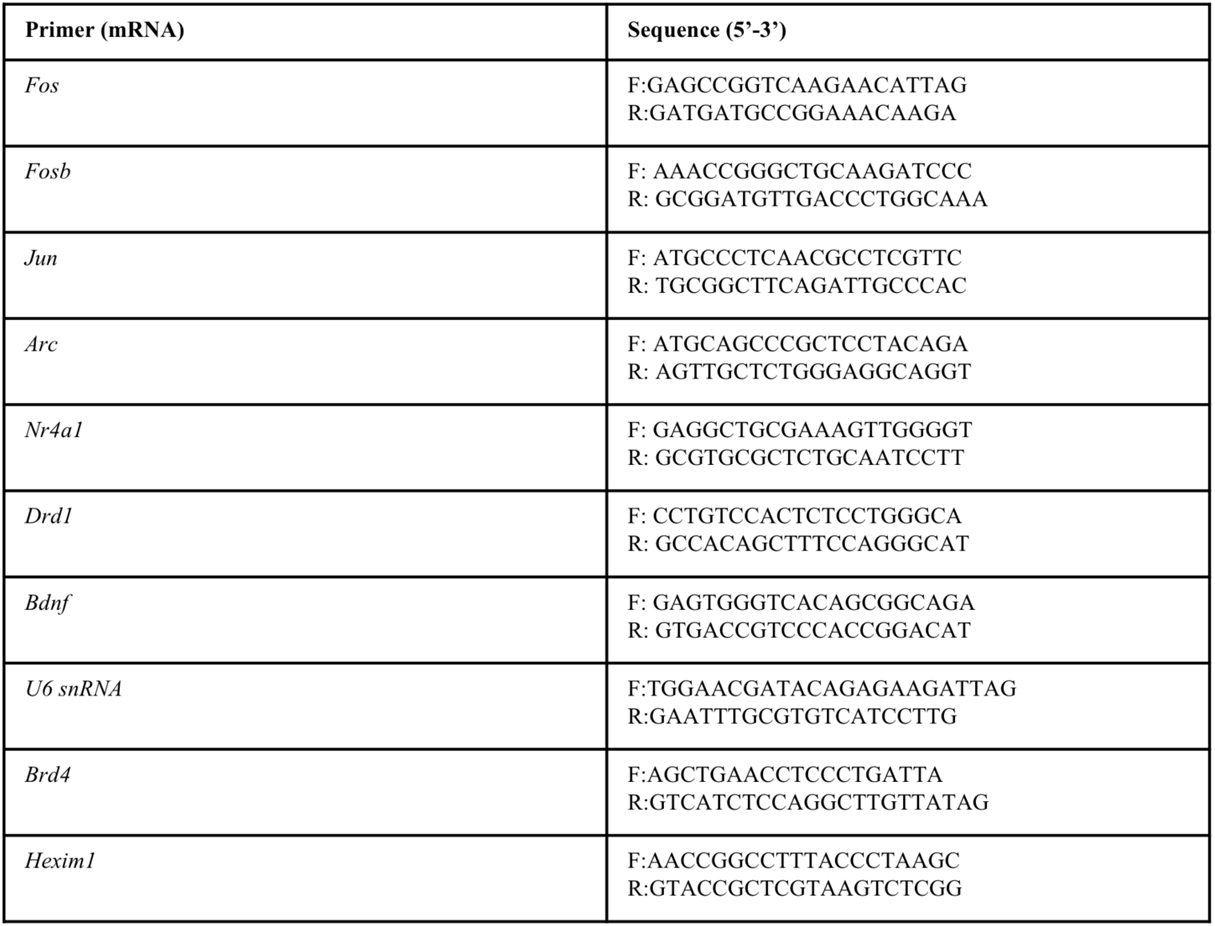

### RNA-seq analysis

For RNA sequencing experiments, primary neurons were plated on 6-well culture dishes (ThermoFisher) at a density of 2 million cells per well and cultured for 7 days prior to drug treatments. RNA was isolated with the RNeasy™ Mini Kit (Qiagen) according to manufacturer’s instructions, and RNA quality assessed by Bioanalyzer (Agilent). Libraries were prepared using the NEBNext™ rRNA-depleted (HMR) stranded library kit and paired-end 100bp sequencing to a depth of 25 million reads per sample was performed on the Illumina NovaSeq™ 6000 at the McGill University and Génome Québec Innovation Centre. RNAseq analysis was performed as we have previously described (45). Quality of reads was determined using FastQC, and trimmed with TrimGalore (0.6.0) (74, 75). Trimmed reads were then aligned to the Ensembl rat reference genome (Rattus_norvegicus.Rnor_6.0.94) (76) with STAR (2.7.1a) (77). Transcripts were assembled with StringTie (1.3.4d) (78) and differential gene expression was assessed with DESeq2 (1.24.0) with the independent hypothesis weighting (IHW) library for multiple testing adjustment (79, 80). Adjusted p value threshold of 0.01 was used for differential expression comparisons. Volcano plots were generated with the EnhancedVolcano (3.13) package. Heatmaps and K-means clustering was completed with pheatmap, and the removeBatchEffect function from limma (3.40.6) (81) was used prior to data visualization. Gene Ontology and gene set enrichment analysis were performed using the WebGestalt browser interface (82) with the Gene Ontology, KEGG and Reactome databases. Significance testing for the overlap of gene lists was performed with the GeneOverlap (1.2.8) package.

### Brd4 chromatin immunoprecipitation (ChIP) and qPCR

For ChIP experiments, 7 million primary neurons were plated on 60 mm culture dishes (ThermoFisher) and cultured for 7 days. Brd4 ChIP was performed as previously described (45). Following treatment, neurons were fixed with 1% formaldehyde in Neurobasal™-A media for 10 minutes at room temperature. Formaldehyde was quenched with glycine added to 125 mM final concentration and then washed once with cold PBS. Cells were placed on ice, collected in cold PBS with 1 mM PMSF and 1x protease inhibitor cocktail (P8340 Sigma) and pelleted at 800 g for 5 min at 4°C. The pellet was resuspended in lysis buffer (10 mM TrisHCl pH 8.0, 10 mM EDTA, 0.5 mM EGTA, 0.25% Triton X-100, 1 mM PMSF, 1x protease inhibitor cocktail) and incubated for 10 min at 4°C on a nutator. Nuclei were pelleted at 800 g for 5 min at 4°C and resuspended in nuclei lysis buffer (50 mM TrisHCl pH 8.0, 10 mM EDTA, 1% SDS, 1 mM PMSF, 1x protease inhibitor cocktail). Nuclei were incubated for 15 min on ice and then sonicated using a BioRuptor (18 cycles, 30 s on/off, high power). Samples were centrifuged at 14 000 g for 10 min at 4°C to remove insoluble debris. A 10% aliquot was taken for chromatin quantification and the remaining sample stored at −80°C until use. The aliquot was incubated at 65°C overnight to reverse crosslinks, treated with RNase A (50 μg/mL) for 15 min at 37°C, and then treated with proteinase K (200 μg/mL) for 1.5 h at 42°C. DNA was isolated and purified by phenol/chloroform extraction followed by precipitation at −80°C with 0.3M sodium acetate pH 5.2, 2.5 volumes of 100% ethanol, and 20 µg of glycogen. DNA was pelleted by centrifugation, washed with 70% ethanol, resuspended in ddH_2_O and quantified using a NanoDrop™ spectrophotometer.

For ChIP, 10 μg of chromatin was used per IP. Chromatin was diluted 9x with dilution buffer (16.7 mM TrisHCl pH 8.0, 1.2 mM EDTA, 167 mM NaCl, 0.01% SDS, 1.1% Triton X-100, 1x protease inhibitor cocktail) For spike-in normalization, *S. pombe* chromatin was prepared as previously described (83), and added to each IP at a constant volume of 2.5 μl. A 1% input sample was removed, and then anti-Brd4 rabbit polyclonal antibody (2 μg, Bethyl, A301-985A) or rabbit IgG antibody (2 μg, Thermo Fisher), as well as anti-*S. pombe* H2B antibody (1 μg, Abcam, ab188271), were added to each IP and incubated at 4°C overnight on a nutator. The following day, 15 μL of Protein G Dynabeads were added and incubation was continued for 4 hours. Beads were washed 2X with low salt buffer (20 mM TrisHCl pH 8.0, 2 mM EDTA, 150 mM NaCl, 0.1% SDS, 1% Triton X-100), 2X with high salt buffer (20 mM TrisHCl pH 8.0, 2 mM EDTA, 500 mM NaCl, 0.1% SDS, 1% Triton X-100), 1X with LiCl buffer (10 mM Tris pH 8.0, 1 mM EDTA, 0.25 M LiCl, 1% NP-40, 1% deoxycholate), 1X with TE buffer (10 mM TrisHCl pH 8.0, 1 mM EDTA) at 4°C. Beads were resuspended in elution buffer (200 mM NaCl, 1% (w/v) SDS) and heated at 65°C for 20 min to elute chromatin. The eluted chromatin was incubated at 65°C overnight to reverse crosslinks and then incubated with proteinase K (200 μg/mL) for 2 h at 37°C. DNA was purified as described above, and resuspended in 200 μL of ddH_2_O for qPCR.

Localization was assessed by qPCR with primers for specific genomic loci; a primer pair amplifying *S. pombe cdc2^+^* was used for spike-in normalization to control of IP efficiency. All qPCR reactions were performed using SSO-advance SYBR Green Supermix (Bio-Rad) and a ViiA 7 Real Time PCR System (Applied Biosystems). For each primer pair in a given experimental condition, percent input for IgG control IP was subtracted from the percent input for the Brd4 IP, followed by normalization to the percent input of *S. pombe cdc2^+^* spike in control. Primer sequences were designed using NCBI Primer Blast. Primers were supplied by IDT and validated for specificity and efficiency before use. Primer sequences for ChIP-qPCR are presented in Table 3.

**Table 3.**
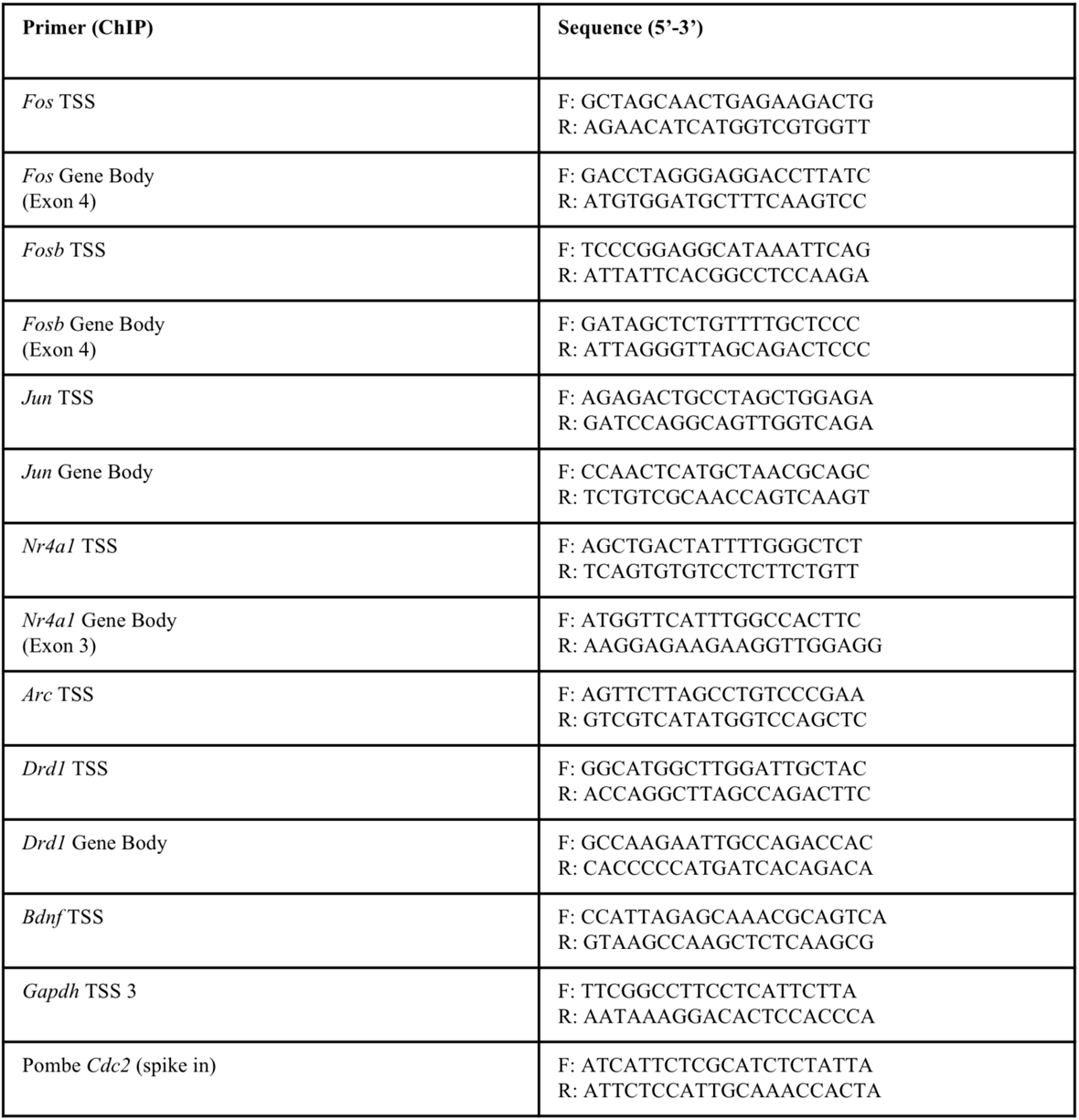

### Protein extraction and western blotting

For determination of Brd4 expression by western blotting, primary neurons were plated on 6-well culture dishes (ThermoFisher) at a density of 2 million cells per well and cultured for 3 days prior to siRNA transfection. 72 hours after transfection, cells were lysed in RIPA buffer (1% NP-40, 50 mM Tris-HCl ph 7.4, 150 mM NaCl, 1 mM EDTA, 1 mM EGTA, 0.1% SDS, 0.5% sodium deoxycholate) and protein was quantified by Bradford assay. Proteins were denatured at 65°C for 15 min in Laemmli buffer and protein expression was assessed by western blot. Western blots were probed with anti-Brd4 (1:2000, Bethyl, A301-985A) or anti-beta-tubulin (1:5000, Life Technology #322600) in 5% milk overnight at 4°C. The following day, blots were visualized with peroxidase-conjugated secondary antibodies and a chemiluminescence detection system.

### Statistical analysis

Statistical testing was performed using GraphPad Prism 9. Biological replicates were defined as a batch of cells derived from a single group of pups (e.g. an n = 6 indicates that distinct 6 batches of pups were used). Primary cells derived from the same group of pups but assigned to different treatment conditions wells were treated as a paired samples for the purpose of statistical analysis. For Figs. 3 and 9, two-way repeated measures ANOVA were performed using gene and siRNA or treatment and pre-treatment as factors respectively. Multiple comparison testing was performed by paired t-tests with Bonferroni correction. For figures where multiple genes were measured (Figs. 2, 3, 4, and 7) Bonferroni correction for multiple comparisons was applied to multiple drug conditions (or in the case of ChIP experiments, multiple locations on the same gene), but not across multiple genes. The threshold for statistical significance was p<0.05 (2-tailed).

## Acknowledgements

This work was supported by grants from the CIHR. J.J.T was supported by fellowships from the CIHR and the McGill Healthy Brains for Healthy Lives initiative. R.M. was supported by fellowships from the McGill-CIHR Drug Development Training Program and the McGill Faculty of Medicine. J.J.C. was supported by fellowships from FRQS. We thank Dr. Michiyuki Matsuda (Kyoto University) for providing us with biosensor constructs and Dr. Hiroyuki Hioki (Kyoto University) for providing AAV plasmids. T.E.H. is the holder of the Canadian Pacific Chair in Biotechnology. We thank the McGill Pharmacology and Therapeutics Imaging and Molecular Biology platform, as well as the McGill Advanced Bioimaging Facility for training and assistance with microscopy and image analysis. Lastly, we thank members of the Hébert, Clarke and Tanny labs for feedback and guidance throughout the development of the project.

## Author Contributions

**J.J.T.**, Conceptualization, Methodology, Investigation, Writing – Original Draft. **R.D.M.**, Conceptualization, Methodology, Software, Writing - Review & Editing. **J.J.C.,** Resources, Writing - Review & Editing **J.C.T.**, Conceptualization, Writing – Review & Editing, Supervision. **P.B.S.C.**, Conceptualization, Writing - Review & Editing, Supervision. **T.E.H.**, Conceptualization, Writing - Review & Editing, Supervision.

## Figure Legends

**Figure S1.**
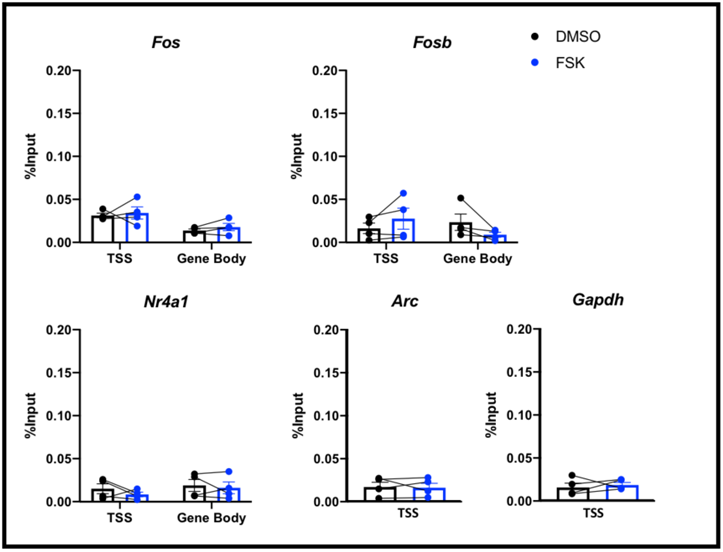
Brd4 is not detectably recruited to chromatin in response to cAMP signaling in cortical neurons. ChIP-qPCR showing Brd4 binding to chromatin in cortical neurons. Values are shown as the percentage input normalized to a spike-in control (see *Experimental Procedures*). Bars depict mean ± SEM, connecting lines indicate biological replicates, n = 4.

**Figure S2.**
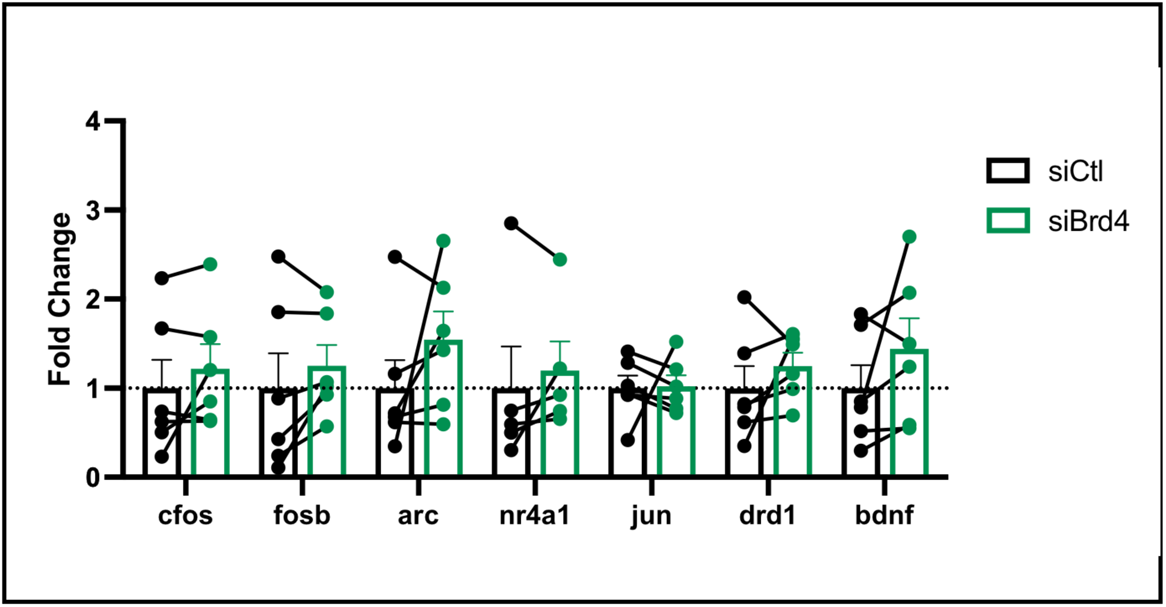
Effect of Brd4 knockdown on baseline gene expression. Relative gene expression 72 hours after siRNA transfection in cells treated for 60 minutes with vehicle (DMSO), determined by RT-qPCR. Points depict biological replicates and bars depict mean ± SEM.

**Figure S3.**
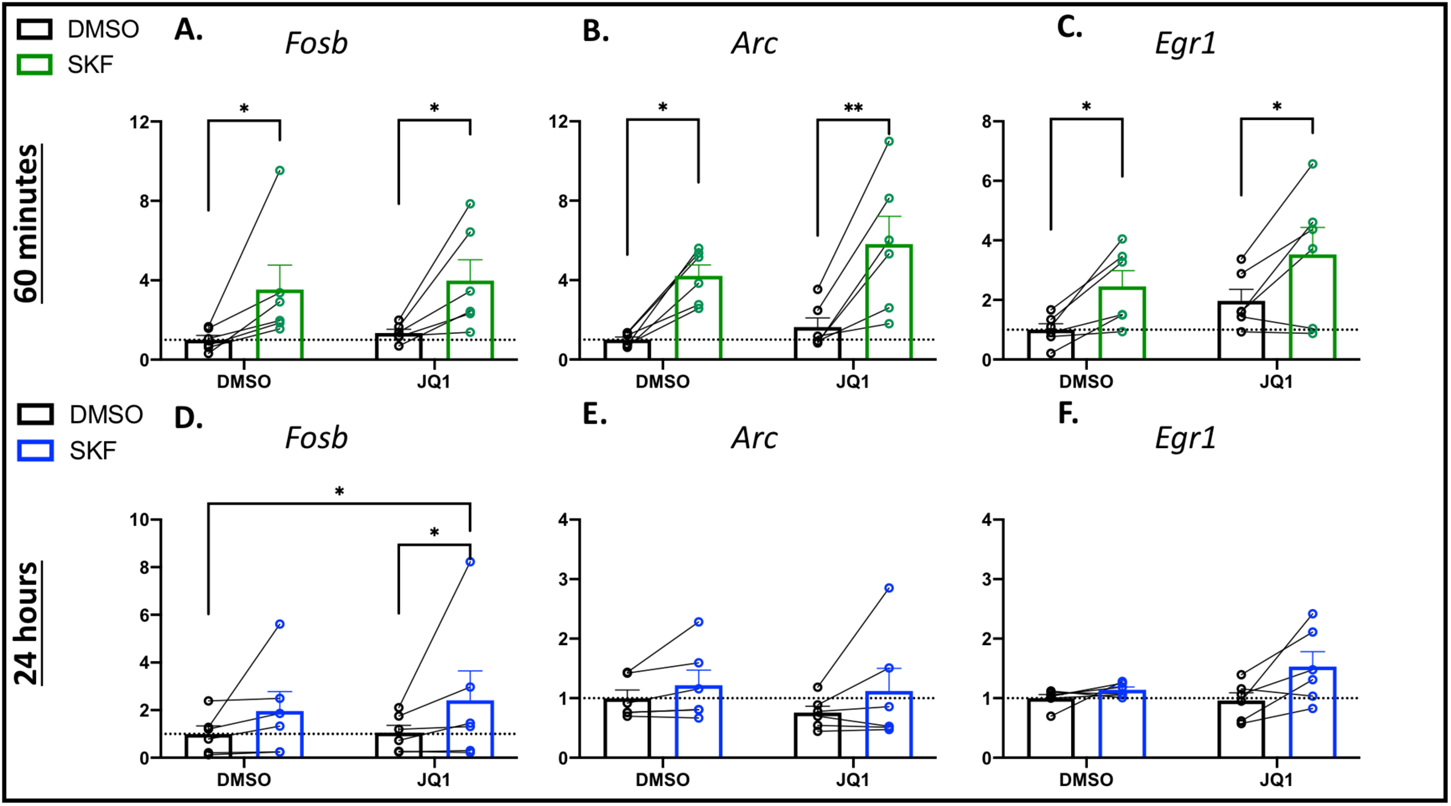
Effect of JQ1 on additional immediate early genes. Expression of three IEGs (*Fosb*, *Arc, Egr1*) was measured by RT-qPCR either 60 minutes (A) or 24 hours (B) after treatment with vehicle (DMSO) or 1 μM of the D1 receptor agonist SKF 81297 (SKF), with or without 2 μM JQ1. Bars depict mean ± SEM, connecting lines indicate biological replicates, n = 6 biological replicates, * p<0.05, ** p<0.01 (Bonferroni corrected paired t-tests, select comparisons shown).

**Figure S4:**
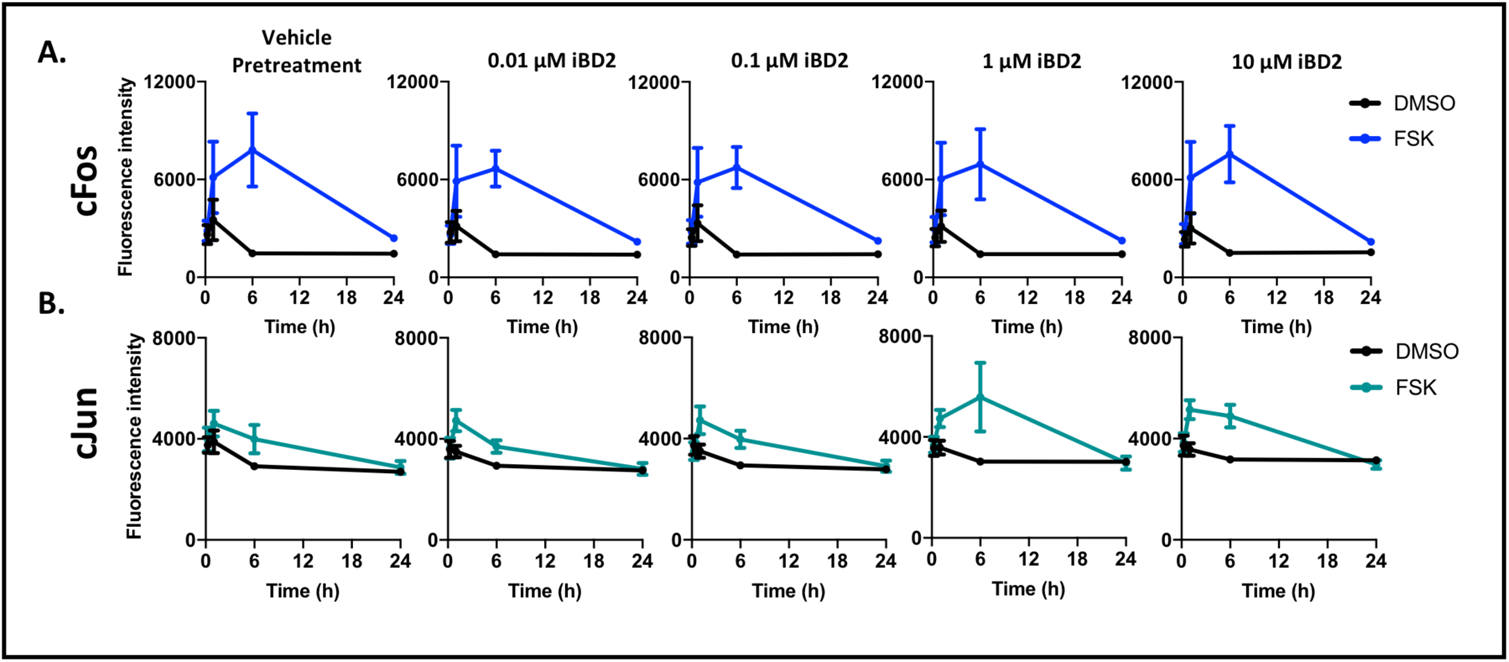
BD2 inhibitor iBD2 does not affect cFos or cJun protein expression. **A,B.** Quantification of cFos (A) and cJun (B) fluorescence intensity in striatal neurons treated with vehicle (DMSO) or 5 μM forskolin (FSK) and iBD2 ranging from 0 to 10 μM. Neurons were treated for 15 minutes, 60 minutes, 6 hours or 24 hours. Points depict mean ± SEM, n = 4 biological replicates.

**Figure S5:**
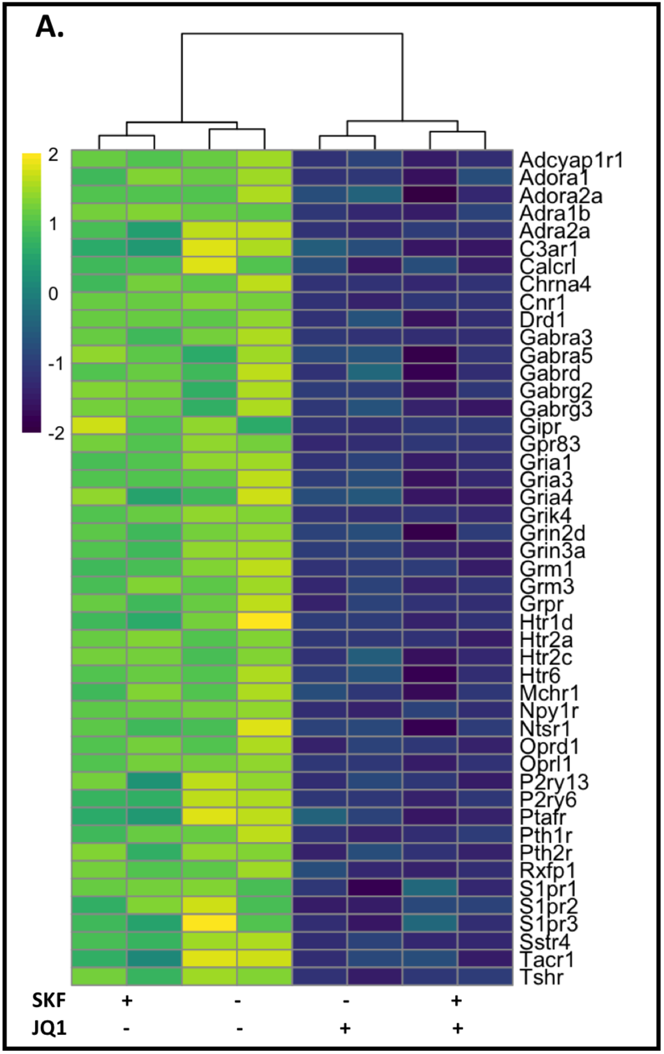
JQ1 reduces expression of GPCRs and ligand-gated ion channels in striatal neurons. **A.** Heat map showing the normalized z score for genes differentially regulated by JQ1 (Log_2_ Fold Change > 0.5, and p < 0.05 for 2 μM JQ1 relative to vehicle/vehicle condition) within the KEGG “neuroactive ligand-receptor interaction” annotation (see Fig. 8C).

**Figure S6:**
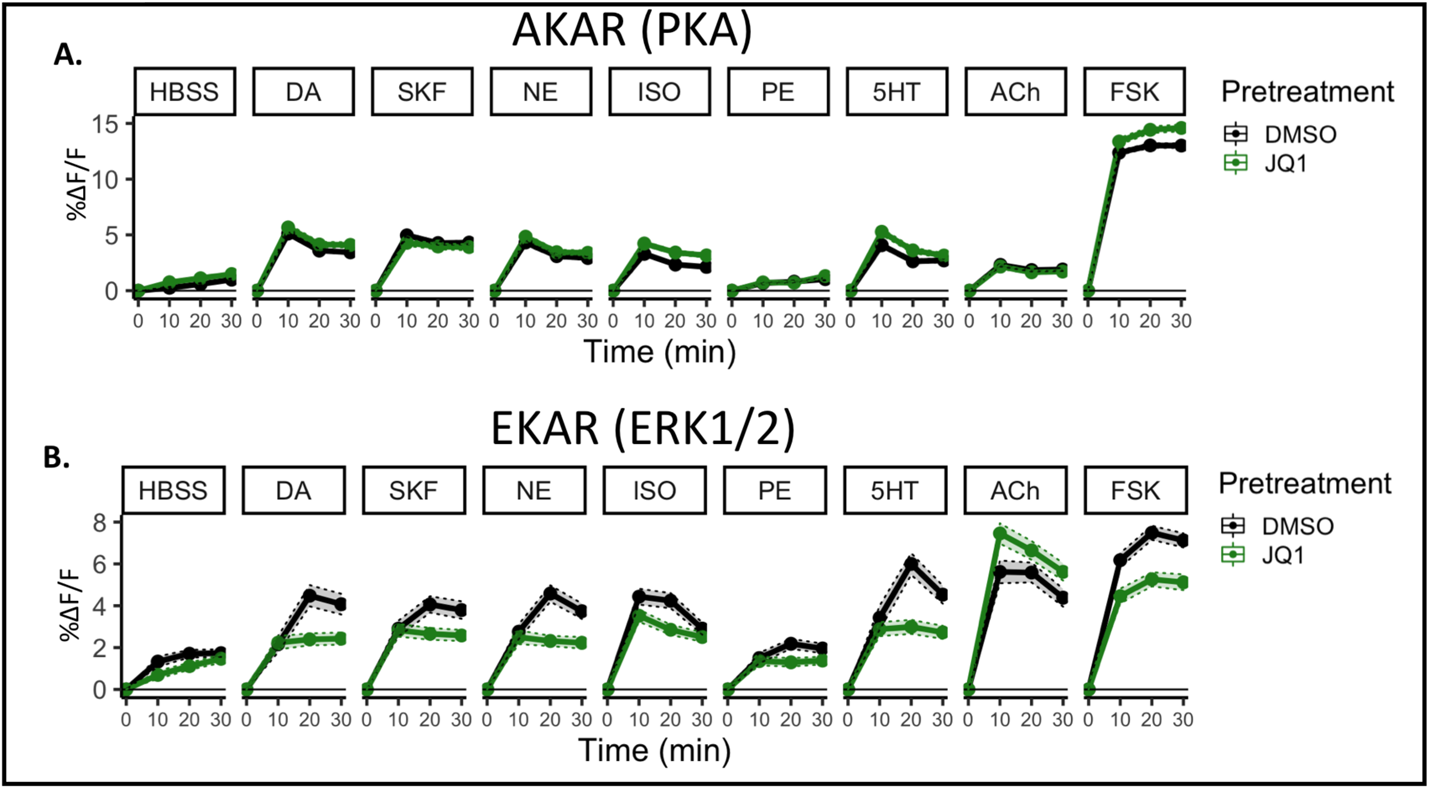
Time course of PKA and ERK1/2 activation in striatal neurons pretreated with vehicle or JQ1. **A.** Time course of PKA activation measured using the AKAR biosensor. PKA activation measured as percent change in FRET ratio (%ΔF/F) after treatment with the indicated ligands. n = 3 biological replicates. **B.** Time course of ERK1/2 activation measured using the EKAR biosensor. ERK1/2 activation measured as percent change in FRET ratio (%ΔF/F) after treatment with the indicated ligands. n = 3 biological replicates.

